# The *SNCA* A53T mutation sensitizes human neurons and microglia to ferroptosis

**DOI:** 10.1101/2025.10.13.682089

**Authors:** Laura Mahoney-Sánchez, Hannah Lucas-Clarke, Alexis Penverne, James R. Evans, Karishma D’Sa, Stephanie Strohbuecker, Patricia Lopez Garcia, Katharina Cosker, Darija Soltic, Ben O’Callaghan, Alexander Griffiths, Sean A. Pintchovski, Helene Plun-Favreau, Jenny Hallqvist, Kevin Mills, Sonia Gandhi

## Abstract

The major pathological hallmarks of sporadic and familial forms of Parkinson’s disease (PD) are the targeted and progressive loss of midbrain dopaminergic neurons (mDA), associated with systemic iron accumulation, α-synuclein (αsyn) accumulation and aggregation, and lipid peroxidation amongst other reactive oxygen species (ROS) generation. Therapeutic strategies aimed towards dopamine restoration, αsyn removal and iron chelation have provided symptomatic relief but failed to prevent or slow disease progression. This is in part due to the lack of understanding of the exact pathways leading to neuronal death in PD. In this study, we investigate ferroptosis, a unique cell death mechanism sharing multiple features with PD pathology, as a relevant pathway with implications in disease pathogenesis. We identified an enrichment of ferroptosis genes dysregulated throughout PD postmortem brain samples and several neuronal and glial PD models. Using CRISPR/Cas9 technology, we generated a rapid iPSC-derived synucleinopathy neuronal model harbouring the *SNCA* A53T mutation and report increased ROS generation, reduced levels of antioxidant glutathione (GSH), impaired mitophagy and a heightened vulnerability to ferroptosis-induced lipid peroxidation and cell death. Critically, inhibition of the key lipid peroxidation enzyme and driver of ferroptosis, 15-lipoxygenase (15-LO), rescued synucleinopathy associated pathologies and prevented pathological αsyn oligomerisation in *SNCA* A53T neurons. Furthermore, we report enhanced microglial ferroptosis susceptibility in models of synucleinopathy. In summary, we highlight a new mechanism by which the familial PD-associated *SNCA* A53T mutation causes cell death and propose 15-LO inhibition as a tractable therapeutic opportunity in PD.

## Introduction

Parkinson’s disease (PD) is the second most prevalent neurodegenerative disease, characterised by the selective and progressive loss of midbrain dopaminergic neurons (mDA) associated with iron accumulation^1^, altered redox state^2,3^ including elevated levels of the lipid peroxidation product 4-hydroxynonenal (4-HNE)^4–6^, neuroinflammation^7^ and α-synuclein (αsyn) aggregation and deposition in the form of Lewy bodies. Mutations in the *SNCA* gene, which encodes αsyn, along with locus multiplications, have been linked to early-onset, rapidly progressive forms of the disease^8–10^. Human-induced pluripotent stem cell (hiPSC)-derived neuronal models carrying the *SNCA* A53T mutation or locus triplication have shed light on several pathogenic pathways in PD, including mitochondrial and lysosomal dysfunction^11,12^, impaired protein and lipid homeostasis^13^, abnormal calcium influx^14^ and excess reactive oxygen species (ROS) generation^15,16^. The critical points of convergence of these cellular phenotypes remain unknown, posing significant challenges for the development of effective disease-modifying strategies to prevent neuronal death.

Ferroptosis is a regulated iron-dependent form of cell death, distinct from apoptosis, that results from the accumulation of lipid hydroperoxides to toxic levels^17^. Whilst the importance of the pathway in normal physiology remains elusive, its involvement in disease pathology has been reported in the context of several cancers and neurodegenerative disorders including Alzheimer’s disease (AD)^18^, amyotrophic lateral sclerosis (ALS)^19^ and PD^20,21^. Ferroptosis is driven by three key mechanisms: *i.* iron dyshomeostasis, *ii.* lipid peroxidation and *iii.* loss of antioxidant defences. Impaired iron flux, storage and distribution in cells can lead to ferrous iron (Fe^2+^) accumulation in the form of labile iron pool (LIP) which in return can trigger lipid peroxidation, and fragmentation to 4-HNE, either enzymatically or non-enzymatically through the Fenton reaction^22,23^. Phospholipids in membrane bilayers are particularly susceptible to oxidation, especially those containing polyunsaturated fatty acids (PUFAs), due to the weak bis-allylic hydrogen bond^24^. Thus, enzymes responsible for enriching cellular membranes with PUFAs and their oxidation, such as acyl-CoA synthetase long chain family member 4 (ACSL4) and 15-lipoxygenase (15-LO) respectively, are positive regulators of ferroptosis^21,22,25^. To counteract lipid peroxidation, cells heavily rely on glutathione peroxidase 4 (GPX4), which reduces lipid peroxides to alcohols using glutathione (GSH) that is produced using cysteine, mainly generated from cystine imported by the glutamate/cystine antiporter (Xc^-^) encoded by *SLC7A11* and *SLC3A2*.

Interestingly, ferroptosis shares several features with PD pathology including iron dyshomeostasis and overload, GPX4 depletion^26,27^, *SLC7A11* downregulation^28^, reduced GSH levels^29^, elevated lipid peroxidation^4–6,30,31^, impaired mitophagy^32,33^, protection from iron chelators^34^ and protection from ACSL4 inhibition^21^. Together, these well-established disease features suggest that ferroptosis may be implicated in the neurodegeneration observed in PD. Furthermore, several PD-associated genes have been shown to modulate pathways that influence ferroptosis sensitivity, for example aldehyde dehydrogenase 1, *ALD1A1*^35^. Taking these converging lines of evidence together, ferroptosis has become more widely accepted as one of the key underlying mechanisms driving PD. However, it remains unclear how best to intervene and modulate this pathway as a therapeutic approach.

In this study we generate a rapid, efficient and inducible hiPSC model of synucleinopathy— *SNCA* A53T harbouring i3neurons (i3Ns)—in which to investigate the role of ferroptosis, its regulation, effectors, and modulation in PD. We report several disease-relevant phenotypes in *SNCA* A53T i3Ns including elevated cytosolic and mitochondrial ROS, reduced GSH and impaired mitophagy. Furthermore, our findings show a direct involvement of the familial PD-associated mutation, *SNCA* A53T, in heightening the sensitivity of both neurons and microglia to ferroptosis effectors and propose 15-LO inhibition as a potential therapeutic target for αsyn-associated pathologies. Finally, we highlight a microglial ferroptotic signature to A53T αsyn monomers and identify *NUPR1* as a potential αsyn-associated stress response gene. This study supports ferroptosis as a key cell death pathway which may drive the pathology of PD, highlighting the potential for ferroptosis-based therapeutic opportunities as disease-modifying therapies.

## Results

### 1. Ferroptosis genes are dysregulated in Parkinson’s disease patients and models

To investigate the putative role of ferroptosis (Figure 1A) in PD, we utilised a manually curated list of 484 ferroptosis-related genes (FRGs) derived from the literature (FerrDB v2, Table S1A)^36^. This dataset includes genes known to promote, halt or indicate ferroptosis. We assessed whether FRGs were enriched among differentially expressed genes (DEG) in publicly available transcriptomic datasets from both PD patient-derived hiPSC models and postmortem PD brain (summarised in Figure 1B and Table S1B), via over-representation analysis (ORA). FRGs were significantly overrepresented in all four datasets (Figure 1Ci) comparing *i.* iPSC-derived mDAs from healthy controls (HC) vs familial-PD individuals carrying either the *SNCA* A53T mutation or *SNCA* triplication (Athauda2024 p<0.0001 z= 22.1) (Figure 1Cii)^16^, *ii.* post-mortem substantia nigra from sporadic PD patients vs HC (Jian2022 p<0.0001 z= 26.1) (Figure 1Ciii)^37^, *iii.* post-mortem brain regions associated with Braak stage 1 vs 6 (Keo2020 p<0.0001 z= 32.8) (Figure 1Civ)^38^ and *iv.* multiple post-mortem brain regions from PD cases at Braak stage 3/4 vs HC (Evans2025 p<0.0001 z= 37) (Figure 1Cv)^39^.

**Figure 1.**
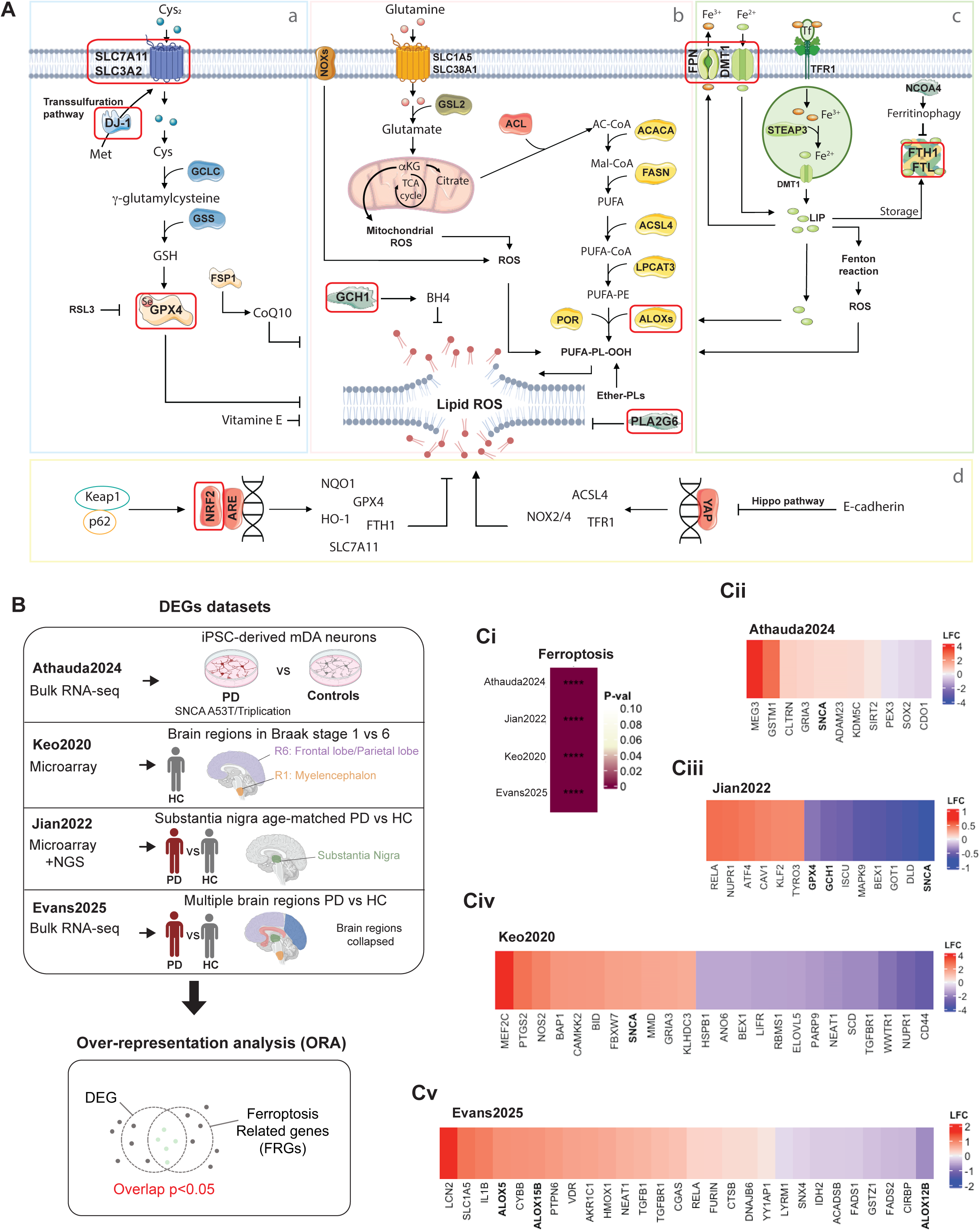
T**r**anscriptomic **evidence for ferroptosis in Parkinson’s disease A.** Schematic of the ferroptosis pathway showing the main regulators: antioxidant machinery (a), lipid metabolism (b), iron metabolism (c) and transcriptional regulators (d). PD associated genes common to the ferroptosis pathway are highlighted in red. **B.** Schematic representation of the publicly available PD transcriptomic datasets and over-representation analysis (ORA) performed in this study. **C.** ORA results highlighting ferroptosis related genes (FRGs) enrichment in the different datasets studied (**Ci**) and the specific genes highlighted from Athauda2024 (**Cii**), Jian2022 (**Ciii**), Keo2020 (**Civ**) and Evans2025 (**Cv**). Schematics made on Biorender.com.

Notably we identified downregulation of key ferroptosis suppressors and antioxidant genes, including *GPX4* and *GCH1,* in the substantia nigra of sporadic PD patients (Figure 1Ciii)^37^. GPX4 is a central regulator of ferroptosis, catalysing the reduction of lipid hydroperoxides (L-OOH) to non-toxic lipid alcohols (L-OH), thereby preventing lipid peroxidation-mediated ferroptosis cell death. The GCH1/tetrahydrobiopterin (BH_4_) axis functions as a master regulator of ferroptosis resistance by supporting endogenous production of antioxidant BH_4_ and modulating CoQ_10_ levels^40^ (Figure 1A). *HMOX1,* which encodes the Heme Oxygenase 1 (HO-1)—an enzyme with dual roles in ferroptosis sensitivity—was significantly upregulated in Braak stage 3/4 brains compared to HC (Figure 1Cv). Furthermore, several members of the lipoxygenase (ALOX) family, which drive PUFA oxidation to promote ferroptosis, (Figure 1A) were also dysregulated in Braak stage 3/4 brains (Figure 1Cv). This included upregulation of *ALOX5* and *ALOX15B,* and down regulation of *ALOX12B.* Additionally, key lipid metabolism genes implicated in determining cellular susceptibility to ferroptosis— *ELOVL5*, *FADS1* and *FADS2—*were found to be differentially expressed in disease (Figure 1Cii-Cv), further supporting that lipid dyshomeostasis and hydroperoxidation contribute to PD pathogenesis.

Collectively, these findings demonstrate that ferroptosis-associated transcriptional alterations are consistently observed across multiple models (iPSC-derived neurons and postmortem brain tissue), disease type (sporadic and familial PD) and Braak stages, underscoring ferroptosis as a potentially critical mechanism in PD progression.

### 2. *SNCA* A53T neurons exhibit oxidative stress phenotypes and increased ferroptosis vulnerability

αsyn aggregation models have previously been shown to exhibit hallmarks of ferroptosis^14,27,41^. In this study, we generated an inducible, genetically editable, and rapid human iPSC-derived neuronal model—i3Neurons (i3Ns)—with an *SNCA* A53T mutation to study ferroptosis. The i3N model system, which enables neuronal differentiation through doxycycline-inducible expression of the transcription factor neurogenin 2 (*NGN2*), as well as knockdown of genes via CRISPRi, has been extensively characterised^42,43^. Using the NGN2/dCas9 iPSC line, we generated stable *SNCA* A53T heterozygous mutant clones via CRIPSR/Cas9 gene editing (Supplementary figure 1F & 1G). A normal karyotype was confirmed for the dCas9 parental line, two CRISPR WT clones, and four of six heterozygous *SNCA* A53T CRISPR clones (Supplementary figure 2). Importantly, the editing process did not impair neuronal differentiation efficiencies, as evidenced by comparable expression of neuronal markers (*MAP2*, *SYN1* and *NeuN*) and a similar downregulation of pluripotency markers *SOX2* and *NANOG* in both WT and A53T i3Ns during differentiation (Supplementary figure 1H & 1I).

Using our rapid synucleinopathy neuronal model, we assessed the generation of cytosolic and mitochondrial ROS (mROS) using the ratiometric dye dihydroethidium (DHE) and fluorescent reporter MitoTracker Red CM-H2XRos, respectively. A53T mutant i3Ns exhibited significantly higher rates of both cytosolic (WT= 1 ± 0.14 vs A53T= 1.47 ± 0.09, p= 0.031) (Figure 2Ai & Aii) and mROS production when compared to WT neurons (WT= 1 ± 0.15 vs A53T= 1.87 ± 0.24, p= 0.039) (Figure 2Bi & Bii). In line with this, levels of lipid hydroperoxidation reducing GSH were decreased in A53T mutant i3Ns as measured by monochlorobimane (mCBl) fluorescence intensity (WT= 1 ± 0.03 vs A53T= 0.89 ± 0.02, p= 0.027) (Figure 2Ci & Cii). These findings confirm that the A53T mutation leads to a pathological oxidative stress environment in our rapid *SNCA* mutant i3N model, consistent with previous observations in PD patient-derived neurons (Virdi et al. 2022; Choi et al. 2022; Athauda et al. 2024).

**Figure 2.**
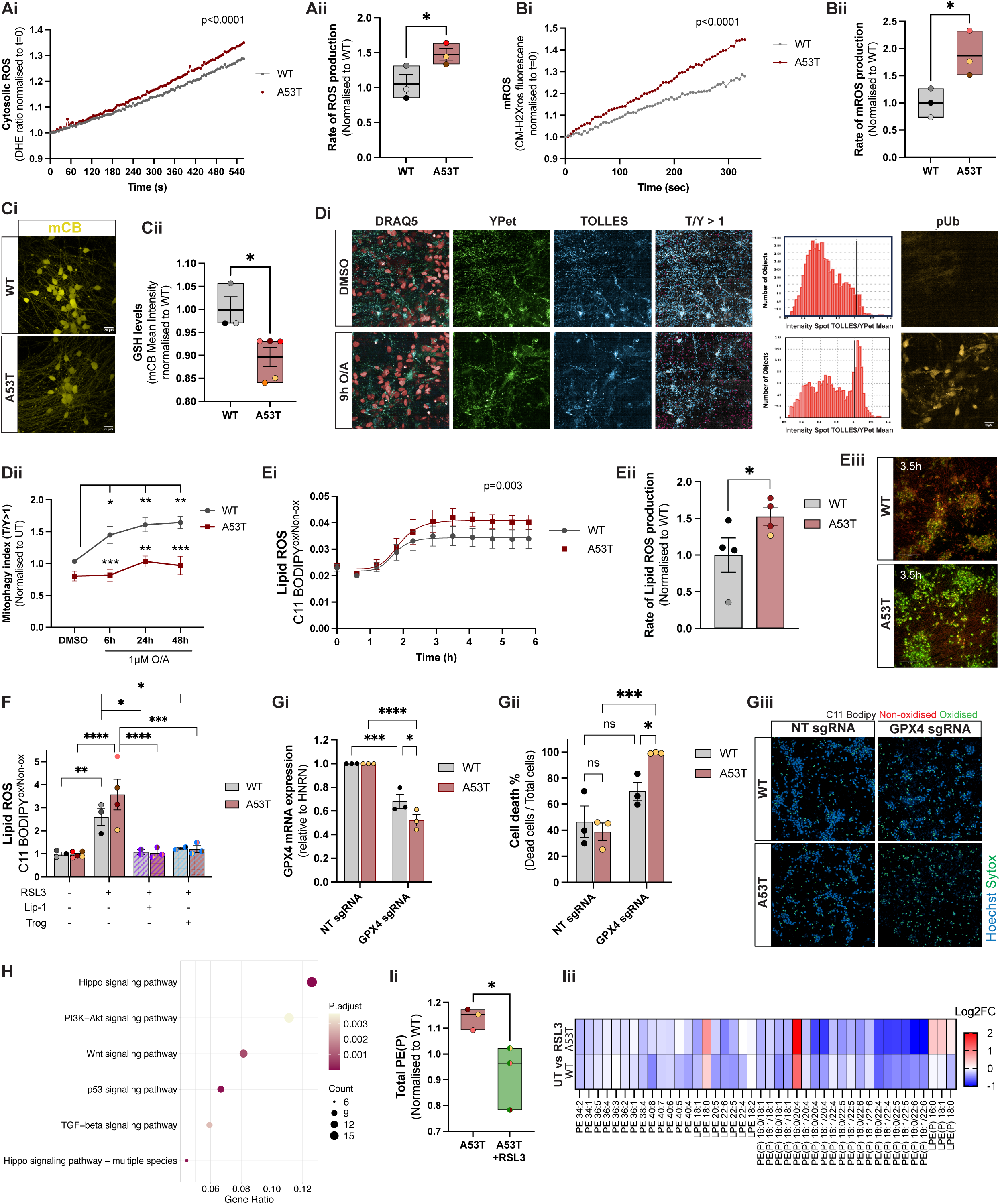
Enhanced oxidative stress and ferroptosis vulnerability in *SNCA* A53T i3neurons. **Ai-Aii.** Trace showing the average ratiometric measurement of basal cytosolic superoxide generation in WT and A53T i3Ns based on dihydroethidium (DHE) ratiometric fluorescence (p< 0.0001, extra sum-of-squares F test) and quantification of the rate of production (*p< 0.05, one-tailed unpaired Student’s t-test). **Bi-Bii.** Trace showing the average basal mitochondrial ROS production in WT and A53T i3Ns based on Mito-CM-H2xROS fluorescence over time (p< 0.0001, extra sum-of-squares F test), and quantification of the rate of production (*p< 0.05, two-tailed unpaired Student’s t-test). **Ci-Cii.** Representative live confocal microscopy images of mCB staining (yellow) and quantification of basal GSH levels in WT and A53T i3Ns as measured by mCB intensity (*p < 0.05, two-tailed unpaired Student’s t-test). **Di.** MitoSRAI i3N neurons treated with 1µM O/A for 9 hours. Mitophagy index is measured by TOLLES /YPet ratio> 1 (blue/green), and pUb antibody staining as a mitophagy initiation marker in i3N neurons. **Dii-Dii.** Time-response of 1µM O/A treatment in WT and A53T i3Ns measuring mitophagy index (*p < 0.05, **p< 0.001, ***p< 0.0001, Two-way ANOVA). **Ei-Eii.** Lipid peroxidation measurements via high-throughput confocal live imaging over 6 hours in WT and A53T i3Ns stained with C11-Bodipy 581/591 and treated with 20µM RSL3 (p< 0.0001, extra sum-of-squares F test), quantification of rate of lipid ROS production (*p<0.005, one-tailed unpaired Student’s t-test), and representative images of i3Ns at 3.5h post-treatment taken via high-throughput confocal live imaging (red non-oxidised, green oxidised). **F.** Lipid peroxidation in WT and A53T i3Ns in response to 20µM RSL3 (3h) +/-co-treatment with 1µM Lip-1 or 2.5µM Troglitazone measured via flow-cytometry (*p< 0.05, **p< 0.001, ***p< 0.0001, Two-way ANOVA). **Gi**. *GPX4* mRNA levels measured via RT-qPCR in WT and A53T i3Ns transduced with either non-targeting or *GPX4* sgRNA lentivirus particles (*p < 0.05, ****p < 0.0001, Two-way ANOVA). **Gii-Giii.** Quantification of percentage of cell death in WT and A53T i3Ns following transduction with either non-targeting or *GPX4* sgRNA lentivirus particles and representative high-throughput confocal live images of i3Ns stained with Hoechst (blue) and SytoxGreen (green). **H.** Dot plot of downregulated KEGG pathways revealed from bulk trancriptomics after 300nM RSL3 treatment in A53T i3Ns. **Ii.** Quantification of total PE(P) species, measured by lipidomics, in A53T i3Ns +/-300nM RSL3 (*p<0.005, two-tailed Student’s t-test). **Iii.** Heatmap of log2 fold change expression of PE species in WT and A53T treated with RSL3 vs untreated. Data shown as mean ± SEM from at least three independent biological replicates or inductions.

To assess mitophagy in neurons, we stably expressed mitoSRAI (signal-retaining autophagy indicator) probe into WT and A53T CRISPRi-i3N iPSC lines^32,44^. Cells were treated with oligomycin (to inhibit ATP synthase/mitochondrial complex V) and antimycin (to inhibit mitochondrial complex III), which in combination (O/A) lead to stress-induced mitophagy. MitoSRAI is a tandem fluorescent protein targeted to the mitochondrial matrix which consists of the yellow fluorescent protein, YPet, fused to the cyan fluorescent protein TOLLES. Unlike YPet, which is readily degraded in lysosomes, TOLLES is resistant to degradation in lysosomal environments. To indicate mitochondrial delivery to lysosomes, we calculated a mitophagy index by segregating mitochondrial units based on the TOLLES:YPet fluorescent intensity ratio. Treatment with 9 hours of O/A results in a shift in the mitophagy index ratio together with an increase in phospho-ubiquitin (pUb), a marker of PINK1 (PTEN-induced kinase 1) dependent mitophagy, suggesting successful induction and detection in i3Ns (Figure 2Di). Interestingly, A53T mutant i3Ns show a significant reduction in O/A-induced mitophagy index compared to WT neurons (WT vs A53T: 6h= 1.45 ± 0.14 vs 0.82 ± 0.091; 24h= 1.6 ± 0.1 vs 1 ± 0.09; 48h= 1.65 ± 0.09 vs 0.97 ± 0.15, p< 0.0001) (Figure 2Dii & 2Diii).

Lipid hydroperoxide accumulation in cellular membranes is considered the central biochemical hallmark and driver of ferroptosis^24,45,46^. To assess lipid peroxidation in WT and A53T mutant neurons, we utilised the ratiometric lipid peroxidation sensor— BODIPY581/591 C11—in the presence of RSL3, a small molecule irreversible inhibitor of GPX4 and established ferroptosis inducer (Figure 1A). The sensitive fluorescent probe integrates into cellular membranes and shifts its emission peak from red (∼590nm) to green (∼510nm) upon lipid membrane hydroperoxidation. Over 6 hours of exposure to RSL3, A53T i3Ns exhibited significantly higher lipid peroxidation levels compared to WT neurons (WT= 1 ± 0.23 vs A53T= 1.53 ± 0.12, p=0.045) _[HLC1]_(Figure 2Ei-Eiii).

Selective ferroptosis induction and higher lipid peroxidation in A53T i3Ns was further confirmed via flow cytometry analysis of BODIPY581/591 C11 in WT and A53T i3Ns in the presence or absence of two well-established ferroptosis blockers: Liproxstatin-1 (Lip-1) and Troglitazone (Trog). Lip-1 acts by directly scavenging lipid peroxides^47^, while Trog, a potent and selective ACSL4 inhibitor^48^, limits PUFA incorporation into membrane phospholipids, thereby reducing lipid hydroperoxidation^25^. Here we observed a significant increase in lipid hydroperoxidation levels in both WT and A53T i3Ns upon a 3 hours RSL3 treatment, with higher levels attained in mutant i3Ns, and a complete reduction to untreated levels with both Lip-1 and Trog (WT: RSL3= 2.6±0.4, RSL3+Lip1= 1.08±0.1, RSL3+Trog= 1.2±0.06, p= 0.01 & p= 0.02; A53T: RSL3 = 3.57 ± 0.7, RSL3+Lip1= 1.03±0.1, RSL3+Trog= 1.21±0.15, p<0.0001 & p=0.0001) (Figure 2F), confirming enhanced vulnerability to RSL3-induced lipid hydroperoxidation in A53T mutant neurons.

We next investigated the effect of reducing *GPX4* levels via CRISPRi in WT and A53T i3Ns. *GPX4* expression was significantly downregulated in both WT and A53T (WT NTsgRNA= 1 vs GPX4sgRNA= 0.68 ± 0.05, p= 0.0003 & A53T NTsgRNA= 1 vs GPX4 sgRNA= 0.52 ± 0.05, p<0.0001) (Figure 2Gi). Importantly, *GPX4* knockdown significantly increased neuronal death with greater toxicity in A53T mutants (WT: NTsgRNA= 46.6 ± 12 vs GPX4sgRNA= 69.7% ± 7.09, p=0.06 and A53T: NTsgRNA= 38.9 ± 6.9 vs GPX4sgRNA= 99.4% ± 0.3, p=0.0006) (Figure 2Gii & Giii).

To investigate the regulation of neuronal ferroptosis, i3Ns were treated with a low dose of RSL3 (300nM) for 6 hours and subjected to bulk-transcriptomics to identify the earliest pathways involved in ferroptosis initiation (Supplementary figure 4A). The effect of RSL3 in WT i3Ns resulted in 11 differentially expressed genes (DEGs) compared to 270 DEGs in the A53T i3Ns (Table S2). KEGG pathway analysis of DEGs after RSL3 treatment in A53T neurons revealed significant downregulation of several transcription factor signalling pathways including Hippo, Wnt, PI3K-Akt, TGFβ and p53 (Figure 2H, Table S3), all of which have been shown to modulate ferroptosis sensitivity via transcriptional regulation^49–52^. The Hippo signalling pathway is a regulatory cascade involved in controlling cell survival, proliferation, and death^53^. Main effectors transcriptional co-activators YAP (Yes-associated protein) and TAZ (transcriptional coactivator with PDZ-binding motif), have been shown to play a dual role in ferroptosis, acting as both promoters and suppressors by regulating transcription of ferroptosis drivers *ACSL4*, *NOX2/4* and *TFR1,* and ferroptosis suppressor *SCL7A11*^49,54,55^. The canonical Wnt pathway involves β-catenin translocation to the nucleus and transcription of target genes such as *GPX4* and *SLC7A11*, thus protecting cells against ferroptosis^50^. In this study, 6 hours exposure to RSL3 in A53T i3Ns resulted in a significant downregulation of the hippo YAP/TAZ transcription factor complex components *YAP1*, *TEAD2* and *TEAD3* and Wnt signalling components including *FZD2/7*, *GNG12*, *LGR5*, *SFRP1/2*, *SOX2*, *TCF7L1*, *WNT5A* and *WNT7B* (Supplementary figure 4B).

In parallel, we performed targeted lipidomic analysis to identify the initial phospholipid changes occurring during neuronal ferroptosis. We focused on the class of phospholipids predominantly affected during ferroptosis-driven lipid peroxidation: phosphatidylethanolamines (PE)^24,56^. RSL3 treatment resulted in a significant and specific reduction in total plasmalogens PE (PE(P)) (A53T= 1.14 ± 0.02 vs A53T+RSL3= 0.92 ± 0.07, p= 0.047) (Figure 2Ii). More specifically we observed a downregulation of PE(P) containing the long-chain polyunsaturated fatty acids known to predominantly oxidise during ferroptosis—arachidonic acid (PE(P) 18:0/20:4 and PE(P) 18:1/20:4) and docosahexaenoic acid (PE(P) 18:0/22:6 and 18:1/22:6) (Figure 2Iii). This reduction in PE(P) was accompanied by a subsequent moderate increase in lysoPE(P) (LPE(P)) supporting the loss of the long fatty acid chain in PE(P) most likely due to their oxidation. Of note, the preferential loss of fatty acids in plasmalogen PE over dyacil PE could be explained by their greater vulnerability to oxidation as a result of the additional ether double bond at the glycerol sn-1 position.

Interestingly, RSL3 did not induce significant changes in PE(P) or LPE(P) in WT neurons, which may be an indication of an absence of lipid peroxidation under low RSL3 concentrations (Table S4).

Finally, we confirmed the pathological lipid peroxidation phenotype and greater ferroptosis vulnerability in patient-derived iPSC cortical neurons harbouring the A53T mutation (Supplementary figure 4C, 4Di & 4Dii), thereby confirming the relevance of ferroptosis in a second, disease-relevant synucleinopathy model. Together, our findings establish a heightened oxidative stress environment and ferroptosis vulnerability in two *SNCA* A53T expressing neuronal models.

### 3. Modulation of lipid peroxidation alters αsyn-driven pathology

Given that the enzymatic oxidation of PUFAs is key to ferroptosis vulnerability in PD models and tissue, we investigated the effect of inhibiting 15-LO—a key lipoxygenase implicated in the oxidation of PUFA in phospholipids during ferroptosis^21,57^ To do this, we assessed the effect of CU-12991, a selective small-molecule 15-LO inhibitor previously characterised in *LRKK2* mutant mDAs^58^, in rescuing pathological phenotypes observed in our rapid pro-oxidant A53T i3N model.

Pre-treatment with CU-12991 significantly reduced the rate of mROS production in A53T i3Ns (A53T= 2.2 ± 0.09 vs A53T + CU-12991= 1.76 ± 0.047, p= 0.0129) (Figure 3Ai & 3Aii), and resulted in a trending decrease in cytosolic ROS (A53T= 1.29 ± 0.14 vs A53T + CU-12991= 0.93 ± 0.29, p= 0.16) (Supplementary figure 5Ai & 5Aii). In parallel, CU-12991 treatment led to a significant increase in intracellular GSH levels in A53T mutant neurons (A53T= 0.96 ± 0.01 vs A53T + CU-12991= 1.12 ± 0.04, p= 0.012), consistent with reduced oxidative burden (Figure 3Bi & 3Bii). Similar rescue effects were also observed in patient iPSC-derived A53T cortical neurons whereby CU-12991 efficiently increased GSH levels (Supplementary figure 5Ci & 5Cii). Collectively, these findings support a role for 15-LO activity in mediating oxidative stress in A53T neuronal models.

**Figure 3.**
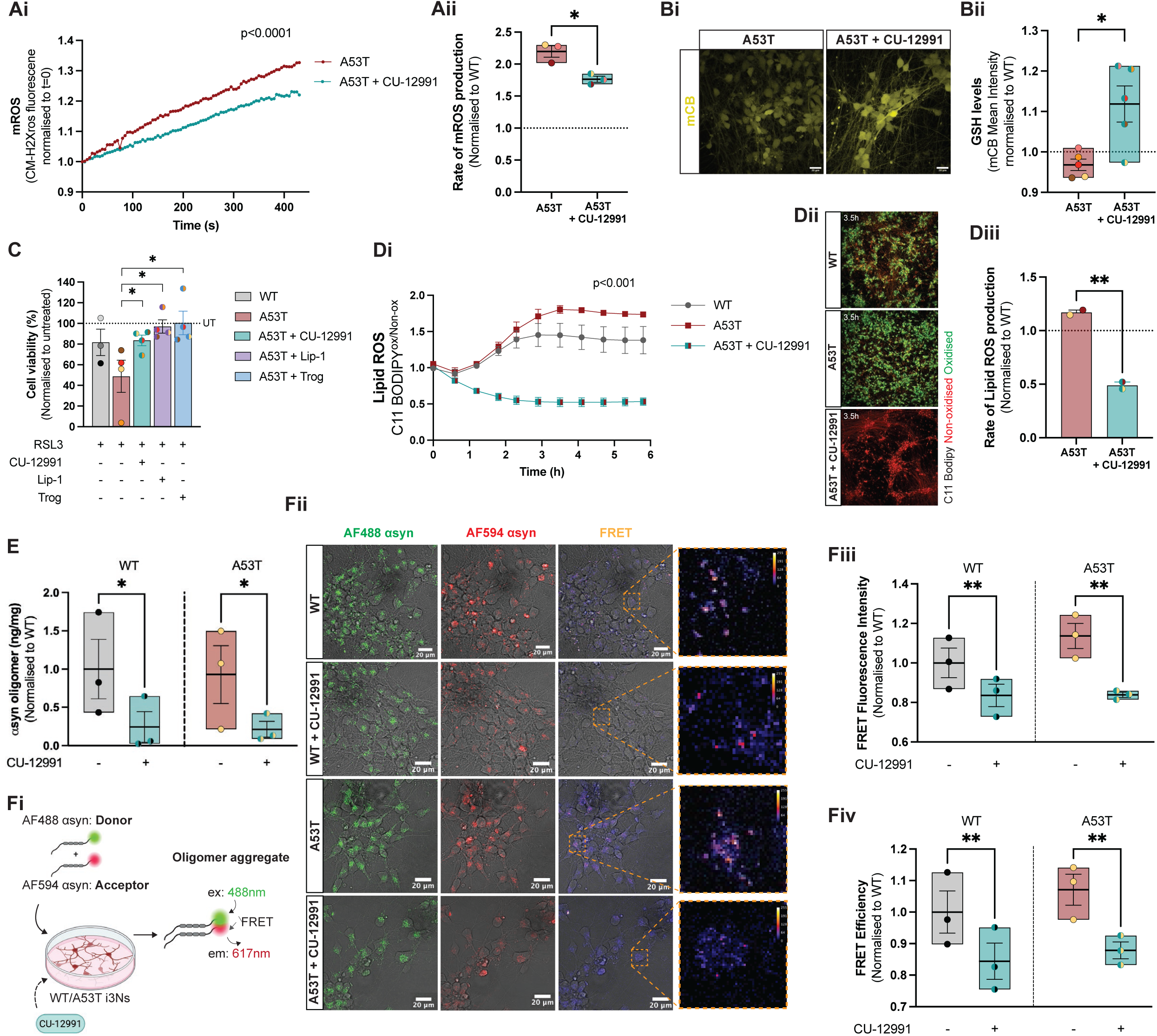
CU-12991 rescues αsyn-associated pathologies in A53T i3neurons **Ai-Aii.** Trace of average mROS production in A53T +/-pre-treatment with 50nM CU-12991 for 5 days based on Mito-CM-H2xROS fluorescence (p< 0.0001, extra sum-of-squares F test) and quantification of mitochondrial ROS production in A53T +/-CU-12991 normalised to WT i3Ns (*p<0.05, two-tailed unpaired Student’s t-test). **Bi-Bii.** Representative live confocal microscopy images of mCB staining (yellow) and quantification of GSH levels in A53T +/-50nM CU-12991 for 5 days as measured by mCB intensity (*p<005, two-tailed unpaired Student’s t-test). **C.** Percentage of cell viability in WT and A53T i3neurons in response to 160nM RSL3 treatment (18h) +/-pre-treatment with 50nM CU-12991, 1µM Lip-1 or 2.5µM Troglitazone (*p < 0.05, one-tailed unpaired Student’s t-test). **Di-Dii.** Lipid peroxidation measurements via high-throughput confocal live imaging over 6 hours in WT and A53T i3neurons treated with 20µM RSL3 +/-pre-treatment with 50nM CU-12991 for 5 days (p < 0.0001, extra sum-of-squares F test), and representative images of i3Ns stained with C11-Bodipy at 3.5h post treatment (red non-oxidised, green oxidised). **Diii.** Quantification of the rate of lipid ROS production in A53T ± CU-12991 (**p= 0.003, two tailed unpaired Student’s t-test). **E.** αsyn oligomer quantification in WT and A53T i3Ns +/-50nM CU-12991 for 5 days normalised to WT UT i3Ns (*p < 0.05, one-tailed unpaired Student’s t-test). **Fi.** Schematic illustration showing how FRET sensor detects aggregation: AF488 αsyn and AF594 αsyn monomers are applied to cells, and the FRET signal is detected within oligomer aggregates. F**ii.** Representative images of AF488 αsyn (green), AF594 αsyn (red) and FRET signals in WT and A53T i3Ns +/-50nM CU-12991 for 5 days. **Fiii-Fiv.** Quantification of FRET fluorescence intensity and FRET efficiency inWT and A53T i3Ns +/-50nM CU-12991 for 5 days (**p<0.0033, **p < 0.0066, two-way ANOVA). Data shown as mean ± SEM from at least three independent biological replicates or inductions. Schematics made on Biorender.com.

Exposure to the ferroptosis inducer RSL3 led to significantly greater neurotoxic effect in A53T i3Ns relative to WT neurons (WT+RSL3= 81.67% ± 12.74 vs A53T+RSL3= 40.19% ± 12.30, p= 0.034) (Figure 3C) supporting our previous finding of heightened ferroptosis susceptibility in A53T mutant neurons. Importantly, pre-treatment with CU-12991 rescued A53T mutant i3Ns from RSL3-induced neurotoxicity to a similar extent as Lip-1 and Trog (A53T+RSL3= 40.2% ± 12.3 vs A53T+CU-12991= 83.58% ± 5.15, p= 0.039) (Figure 3C) and abolished lipid hydroperoxide accumulation (A53T= 1.17 ± 0.02 vs A53T + CU-12991= 0.49 ± 0.03, p= 0.003) (Figure 3Di, 3Dii & 3Diii).

mROS generation and lipid peroxidation are both known to accelerate αsyn oligomerisation and pathological aggregation in neurons^13^. Given the efficiency of CU-12991 in halting oxidative stress in A53T mutant neurons, we next investigated whether 15-LO inhibition could ultimately mitigate the *de novo* formation of toxic αsyn oligomers, which have previously been shown to preferentially form on mitochondrial membrane surfaces under oxidative stress conditions^13,21,59^. Critically, pre-treatment with CU-12991 significanlty reduced αsyn oligomer load in both WT and A53T neurons (WT= 1 ± 0.39 vs WT+CU-12991= 0.24 ± 0.2 & A53T= 0.929 ± 0.377 vs A53T+CU-12991= 0.213 ± 0105, p= 0.026) (Figure 3E). The effect of 15-LO inhibition in preventin αsyn oligomerisation was further assessed using a Förster resonance energy transfer (FRET) pair of fluorescently labelled αsyn monomers. When the fluorophores are in close proximity (<10nm), as is the case within oligomers, energy is transferred from donor to acceptor (FRET signal, Figure 3Fi). Following a 48h incubation with fluorescently labelled αsyn monomers, FRET singal puncta were detected intracellularly in both WT and A53T neurons, indicating aggregate formation (Figure 3Fii). Pre-treatment with CU-12991 significantly reduced FRET Intensity (WT: UT= 1 ± 0.07 vs CU-12991= 0.84 ± 0.06, p= 0.0033 & A53T: UT= 1.14 ±0.06 vs CU-12991= 0.838±0.013, p= 0.033) (Figure 3Fii & 3Fiii), and FRET efficiency in both cell lines (WT: UT= 1 ± 0.07 vs CU-12991= 0.84 ± 0.06 & A53T: UT= 1.07 ± 0.05 vs A53T+CU-12991= 0.88 ± 0.03, p= 0.0066) (Figure 3Fiv & Supplementary figure 5D), suggesting a reduction in αsyn oligomer formation.

In summary, our findings demonstrate that selective inhibition of 15-LO with CU-12991 significantly counters multiple disease-associated phenotypes and protects A53T mutant neurons against ferroptosis-induced lipid peroxidation and cell death.

### 4. *SNCA* A53T mutation increases microglial vulnerability to ferroptosis

Having assessed ferroptosis regulation in synucleinopathy neuronal models, we next explored whether glial cell types exhibit differential susceptibility to ferroptosis in PD. We utilised a publicly available single-cell transcriptomic dataset of the SNpc from 15 sporadic PD and 14 control brains^60^ to investigate FRG enrichment in different cell types (Figure 4A). We found that FRGs were significantly overrepresented in PD-associated microglia (z= 15.6, p<0.0001) and oligodendrocytes (z= 18.5, p<0.0001), while no overlap was detected in astrocytes (Figure 4Bi). Within microglia, the ferroptosis driver genes *NDRG1* and *SMAD7* were upregulated, whereas the suppressor genes *MAPKAP1* and *ACSL3* were downregulated.

**Figure 4.**
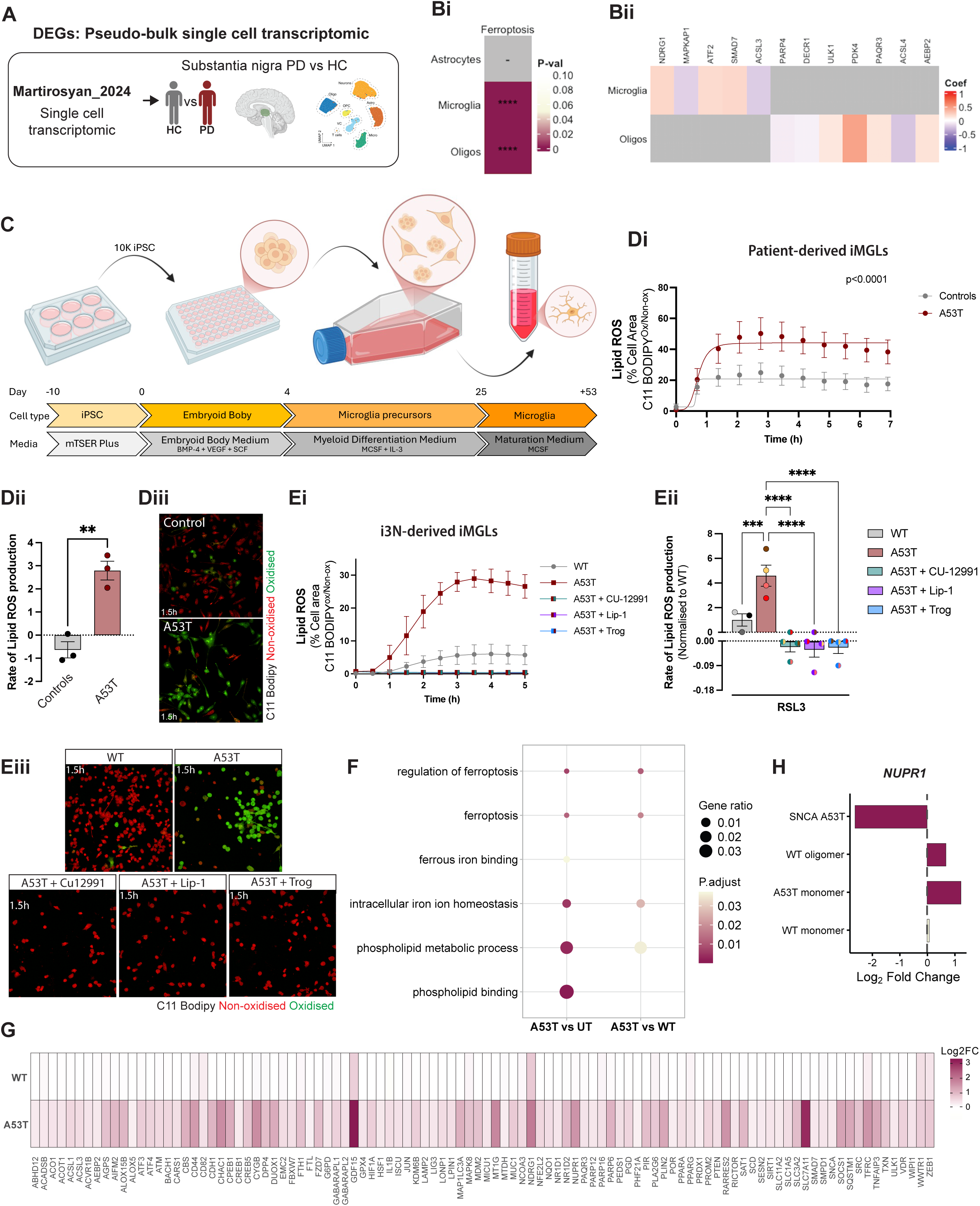
A53T αsyn modulate ferroptosis in microglia **A.** Schematic representation of the publicly available PD single cell transcriptomic dataset used to performed ORA with ferroptosis related genes (FRGs). **B.** ORA results highlighting FRGs are enriched in PD microglia and oligodendrocytes and the specific FRGs differentially expressed in both cell types. **C.** Schematic representation of the protocol used to differentiate human iPSM into microglia like cells (iMGLs). **Di-Diii.** Traces and rates of lipid peroxidation measurements via high-throughput confocal live imaging over 6 hours in A53T patient-derived iMGLs and isogenic controls at ∼day 45 of differentiation treated with 0.5µM RSL3, and representative images of C11-BODIPY (red non-oxidised, green oxidised) stained patient iPSC-derived iMGLs at 1.5h post RSL3 treatment (0.5µM) (p < 0.0001 extra sum-of-squares F test, p=0.003 two-tailed unpaired t-test for rate). **Ei-Eiii.** Traces and rates lipid peroxidation measurements via high-throughput confocal live imaging over 6 hours in i3N WT and A53T CRISPR/Cas9 iPSC-derived iMGLs at ∼day 45 of differentiation treated with 0.5µM RSL3 ± 50nM CU-12991 or 1µM Lip-1 or 2.5µM Trog, and representative images of C11-BODIPY stained iMGLs at 1.5h post RSL3 treatment (0.5µM) (p < 0.0001, extra sum-of-squares F test, (***p<0.001, ****p<0.0001, one-way ANOVA with Dunnett’s multiple comparisons). **F.** Dot plot of dysregulated pathways in healthy control iMGLs untreated vs A53T αsyn monomers and WT vs A53T αsyn monomers from Evans et al. (2025). **G.** Heatmap of log2 fold change expression of FRGs differentially expressed in control iMGLs treated with either WT or A53T αsyn monomers from Evans et al. (2025). **H.** *NUPR1* expression across the iMGL DEGs data set from Evans et al. (2025), showing downregulation in *SNCA* A53T iMGLs and upregulation in response to WT oligomers or A53T monomers.

Notably, the stress-responsive transcription factor *ATF2—*a potential surrogate of NRF2 pathway activation^61^—was significantly upregulated, possibly reflecting a compensatory antioxidant response in PD microglia (Figure 4Bii). In oligodendrocytes, we observed dysregulation of the lipid peroxidation driver gene *ACSL4,* alongside altered expression of *PARP4, DECR1*, and upregulation of multiple ferroptosis drivers including *ULK1, PAQR3* and *AEBP2* (Figure 4Bii). Collectively, these findings suggest that ferroptosis may represent a relevant glia-mediated pathogenic mechanism in PD, with microglia and oligodendrocytes displaying distinct vulnerability signatures (Table S5).

Next, we generated ipsc-derived microglia-like cells (iMGLs) following a previously published protocol (Figure 4C) ^39,62^. Both CRISPR-edited A53T and WT iMGLs successfully differentiated, with confirmed expression of canonical microglial markers IBA1, CX3CR1 and P2YR12 (Supplementary figure 5A & 5B). Phagocytic activity was validated in both genotypes, confirming acquisition of microglial functional phenotypes (Supplementary figure 5C).

Importantly, patient-derived iMGLs harbouring the A53T mutation exhibited a significantly elevated rate of lipid peroxidation following treatment with RSL3, compared to isogenic control iMGLs, as measured by the live-imaging sensor BODIPY581/591 C11 (Isogenic controls=-0.64 ± 0.035 vs A53T= 2.79 ± 0.41, p= 0.003) (Figure 4Di-4Diii). Similar results were observed in iMGLs generated from the CRISPR-edited A53T i3N iPSC cell line, where the introduction of the A53T mutation led to an increased rate of lipid peroxidation in response to RSL3 exposure (WT= 1 ± 0.5 vs A53T= 4.59 ± 0.86, p= 0.0003) (Figure 4Ei-4Eiii). Of note, treatment with ferroptosis inhibitors Lip-1 and Trog completely abolished lipid hydroperoxides accumulation in A53T iMGLs, confirming selective ferroptosis induction (A53T= 4.59 ± 0.86 vs A53T+Lip-1=-0.031 ± 0.027, A53T+Trog=-0.023 ± 0.022, all p<0.0001). Importantly, 15-LO inhibition via CU-12991 produced similar protective effects and prevented lipid peroxidation in A53T mutant iMGLs (A53T= 4.59 ± 0.86 vs A53T+CU-12991=-0.021 ± 0.018, p<0.0001) (Figure 4Ei-4Eiii). Of note, there was no difference in RSL3-induced lipid ROS levels between healthy and *SNCA* A53T mutant patient hiPSC-derived astrocytes suggesting a cell type specific A53T-driven vulnerability to ferroptosis (Supplementary figure 5D).

To assess whether a microglial ferroptosis transcriptomic signature is associated with αsyn aggregation, we utilised our iMGL datasets published in Evans et al. (2025)^39^. Examination of the pathway analysis of the A53T monomeric condition compared to WT monomers or the untreated condition shows enrichment of ferroptosis-associated pathways, including intracellular iron ion homeostasis, regulation of ferroptosis, ferrous iron binding, and ferroptosis itself, alongside phospholipid metabolic process and phospholipid binding pathways (Figure 4F, Supplementary figure 5Ei-Eiii). DEGs from either the WT or A53T monomer treatment with log2 fold change > 0.25 were taken from Evans et al. (2025) and compared against the FRGs list, identifying 102 overlapping genes (Figure 4G). Among the significantly upregulated FRGs were those directly involved in iron import and storage (*TFRC*, *FTH1* and *FTL*), ferritinophay (*NCOA4*), and regulation of cellular iron levels (*IREB2* and *HAMP*) (Figure 4G), in line with dysregulated iron handling and cellular iron overload, which would ultimately trigger ferroptosis. Critically, genes involved in lipid peroxidation and detoxification (*ALOX15B, ALOX5* and GPX4) were equally upregulated in response to A53T αsyn monomer treatment, further suggesting the occurrence of ferroptosis. In parallel, A53T αsyn monomer treatment was also shown to lead to upregulation of several antioxidant defence genes, including *GPX4, NFE2L2* (encoding NRF2), *NQO1, SLC7A11* and *AIFM2 (*also known as ferroptosis suppressor protein 1 – FSP1) (Figure 4G). This coordinated induction of antioxidant responses suggests microglia may attempt to counteract A53T αsyn-induced ferroptotic stress. Further examination of the DEGs across all iMGL PD models (WT monomer, A53T monomer, WT oligomer, and patient-derived *SNCA* A53T mutation)^39^, revealed consistent dysregulation of the stress-induced transcription factor *NUPR1* (Figure 4H). Specifically, *NUPR1* was significantly downregulated in patient-derived *SNCA* A53T iMGLs, whereas both WT oligomers and A53T monomers in control iMGL induced its significant upregulation (Figure 4H). Notably, *NUPR1* was also identified as a significant upregulated gene in our overlap analysis between DEGs from PD vs HC substantia nigra and FRGs (Figure 1Ciii). Interestingly, *NUPR1* has recently been reported as a critical repressor of ferroptosis^63,64^, suggesting it may also play a role in regulating A53T αsyn-mediated microglial ferroptosis.

In summary, our data demonstrate a heightened and cell-type specific vulnerability to ferroptosis in *SNCA* A53T mutant microglia and disruption in iron handling and ferroptosis associated pathways in response to mutant αsyn monomers, suggesting a pathogenic feedback loop. Finally, *NUPR1* emerges as a stress-response gene dysregulated in *SNCA* A53T iMGLs which may play a role in modulating ferroptosis in microglia, revealing a previously unrecognized layer of transcriptional regulation in microglial ferroptotic stress responses.

## Discussion

Parkinson’s disease is a progressive neurodegenerative disorder with a predicted rise in incidence^65^, for which there are no disease-modifying therapies. While the loss of dopaminergic neurons is the main hallmark of PD, the molecular mechanisms driving this selective vulnerability remain unclear. Here, we provide converging evidence that ferroptosis—a regulated, iron dependent form of lipid-peroxidation cell death—may be implicated in PD pathogenesis, particularly in the context of the familial PD *SNCA* A53T mutation.

Through bioinformatic ORA, we revealed enrichment of ferroptosis related genes across PD models and post-mortem brain tissues. Dysregulated expression of genes involved in PUFA metabolism and lipid peroxidation (i.e. *PTGS2, FADS1, ELOVL5, GCH1* and members of the ALOX family) strengthen the implication of ferroptosis-driven lipid peroxidation in PD. Notably, altered expression of *GPX4*, the central enzyme involved in detoxifying ferroptosis-driven lipid hydroperoxides^25^, was identified in PD post-mortem brain tissue, underscoring the disruption of intracellular lipid detoxifying defences. These transcriptomic findings align with previously reported genetic evidence linking PD susceptibility genes and ferroptosis. For instance, mutations in DJ-1, causative of early-onset PD^66^, have been shown to impair GSH biosynthesis and heighten ferroptosis vulnerability^67^. Loss-of-function mutations in *PLA2G6* lead to neurodegeneration with brain iron accumulation (NBIA) disorders with Parkinsonism^68^. The *PLA2G6* gene encodes phospholipase A2 group Vi, an enzyme that has been shown to remove toxic 15-hydroperoxyeicosatetraenoic (15-HpETE), a direct product of 15-LO activity, from phosphatidylethanolamine (PE), thus preventing ferroptosis induction^69^. *GCH1* has been identified as a risk loci in PD genome wide association studies^70^, and loss-of-function mutations in *GCH1* are associated with increased risk of dopa-responsive dystonia and PD^71^. Intriguingly, a recent study has identified the GCH1-tetrahydrobiopterin (BH4)-phospholipid axis as a master regulator of ferroptosis resistance^40^. In addition, hypermethylation in the promoter region of *SLC7A11* in sporadic PD patients was shown to result in its loss-of-function, drop in GSH production, disruption of GPX4 function and subsequently increase in susceptibility to ferroptosis^28^.

In this study, we examined cellular pathology in the context of the PD autosomal-dominant *SNCA* A53T mutation. We generated a rapid and robust disease relevant synucleinopathy model by introducing the A53T point mutation within the *SNCA* gene of the inducible NGN2 iPSC line (i3N). These neurons recapitulated key pathological traits previously observed in patient-derived A53T cortical and mDA neurons^13,15^, including elevated cytosolic and mitochondrial ROS, as well as reduction of GSH levels. Here, we further established a neuronal model for real-time detection of mitophagy, a process recognised as one of the central mechanisms in PD pathogenesis which until now, has proved challenging to detect in neurons^72^. Critically, we showed that mitophagy is decreased in A53T neurons compared to WT, further strengthening the link between αsyn and mitophagy defects in PD. Interestingly, a recent study provided direct evidence that efficient mitophagy protects cells against ferroptosis by reducing mROS^73^. This suggests that the increased mROS levels and ferroptosis vulnerability observed in A53T neurons may result from their impaired ability to selectively degrade damaged mitochondria.

This model has enabled us to dissect early ferroptosis signalling events and phospholipid changes, and demonstrates the heightened vulnerability conferred by the A53T mutation, further implicating this cell death pathway in familial PD forms. In agreement with our findings, a recent study showed increased lipid peroxidation markers, iron accumulation, and reduced GSH in *SNCA* A53T transgenic mice models, and enhanced vulnerability to RSL3 induced cell death and lipid peroxidation in A53T overexpressing cells^27^. Other studies have shown that αsyn levels modulate ferroptosis sensitivity in iPSCs-derived midbrain neurons whereby reducing αsyn protected neurons against ferroptosis while overexpression associated with *SNCA* triplication renders mDAs more vulnerable to ferroptosis^41^.

Our targeted phospholipidomic profiling revealed PE(P) and LPE(P) as the class of phospholipids primarily dysregulated in A53T neurons under a ferroptosis stressor, pointing towards the preferential oxidation of PE(P) in A53T neurons. This class of phospholipids, enriched in neuronal membranes and characterised by heightened oxidative liability due to an additional double bond, likely serve as a key pool of PUFA available for peroxidation and ferroptosis execution^74,75^. This theory is supported by a recent study suggesting that ferroptosis unfolding in neurons may be governed by peroxidation of PE(P)^41^. Intriguingly, PE(P) species have also been reported to be decreased in plasma from AD and PD patients, raising the possibility of peripheral biomarkers reflective of central lipid peroxidation^76^.

The A53T mutation in αsyn is located within the lipid binding domain and is thought to increase lipid-protein binding^13^. Furthermore, mitochondrial cardiolipin was reported to increase αsyn oligomerisation, disruption of the membrane and production of mROS^13,14^. Therefore, a potential mechanism by which A53T increases neuronal ferroptosis is by increased binding of αsyn to membrane lipids, ROS generation, and subsequent lipid peroxidation driven ferroptosis cell death. Conversely, ferroptosis may in turn increase αsyn oligomerisation as membrane lipid peroxidation products are known to drive and stabilise αsyn oligomerisation^77^, therefore creating a vicious toxic loop which may be contributing to synucleinopathy associated pathogenesis. Interestingly, examining pathway analysis from iMGL DEGs datasets recently published in (Evans et al. 2025), highlight upregulation of phospholipid metabolic process and phospholipid binding in iMGLs exposed to A53T αsyn monomers. Determining the dynamics of the relationship between PUFAs and A53T αsyn will be vital in deciphering this mechanism. Here we show that blocking one of the main drivers of enzymatic phospholipid hydroperoxidation, 15-LO, rescued A53T associated oxidative stress pathologies including mitochondrial ROS production and elevated intracellular GSH levels. Critically, restoring the redox balance ultimately resulted in reduced αsyn oligomerisation, positioning 15-LO as a promising therapeutic target in PD. Interestingly, a recent study has shown further therapeutic benefits of 15-LO inhibition via CU-12991 in preventing 4-HNE post-translational modifications of LRRK2 kinase and its subsequent pathogenic hyperactivation^58^.

Investigating ferroptosis in PD patient-derived microglia highlight iron dysregulation as a potential central driver of ferroptosis in relation to the *SNCA* A53T mutation. Analysis of our iMGL bulk RNA sequencing datasets^39^ revealed that exposure to A53T αsyn monomers, but not WT monomers, led to dysregulation in multiple iron regulatory genes accompanied by the concurrent upregulation of antioxidants genes. While microglia play a central role in protecting neurons against stressors, with age and disease duration, microglia adopt a toxic, disease associated state contributing towards non-cell autonomous neuronal death^78,79^. We suggest that microglial ferroptosis driven by A53T αsyn might initiate a toxic signalling cascade contributing to neuronal death in neurodegenerative disease. This proposition aligns with recent studies showing that sub-lethal microglial ferroptotic stress trigger conversion of astrocytes to a neurotoxic state and culminate in neuronal death^80^. Finally, the dysregulation of *NUPR1*, and the difference in its direction of effect across PD microglial models, suggests that its modulation may depend on the cellular stress state. Specifically, A53T αsyn monomers and WT αsyn oligomers initiate an acute stress and immune activation within 24 hours, whereas the endogenous *SNCA* A53T mutation is associated with a prolonged, chronic stress and immune state^39^. Interestingly, a recent meta-analysis of PD gene expression identified dysregulated expression of *NUPR1* in PD across three studies and emerged as positively correlated with the immune microenvironment of PD patients^81^. The molecular pathways involving *NUPR1* in PD-associated microglial dysfunction and its relation to synucleinopathy and ferroptosis remains unanswered and provides a prospect for future research.

Taken together, our study provides unique insights into the complexity of CNS ferroptosis and highlights its relevance to PD pathology and more specifically to synucleinopathy. Our approach combining disease-relevant human patient-derived neuronal and microglial models, and targeted transcriptomic analysis, revealed a direct link between the familial PD *SNCA* A53T mutation and heightened susceptibility to ferroptosis, providing novel insights into the mechanisms driving PD pathology. Our research underscored the potential therapeutic benefit of targeting this pathway via 15-LO inhibition, with marked effects in reducing oxidative stress and αsyn pathological oligomerisation, strengthening the notion that 15-LO inhibition has the potential to be an effective disease-modifying therapy for PD.

## Methods

### HEK293 culture and sgRNA lentivirus production

HEK293 cells were cultured in DMEM (Gibco, cat. no. 31966021) supplemented with 10% FBS, 1% Glutamax (cat. no. 35050038) and 1% penicillin-streptomycin (cat. no. 15070063). For CRISPRi knockdown, sgRNAs were packaged into lentivirus by transfecting HEK293 cells using FuGene HD (Promega, cat. no. E2311) with 3µg of transfer plasmid, 2µg of pxPAX2 (Addgene, #12260) and 1µg of pMD2.G (Addgene, #12259). 18 hours post-transfection, media was replaced with fresh HEK293 media, and the lentivirus particles were harvested 48 hours later by collecting the media and ultracentrifuging it for 3 hours at 48000rcf at 4°C. The lentivirus pellet was resuspended in ice-cold sterile PBS, snap frozen, and aliquoted for future use.

### Human iPSC culture and neuronal differentiation

Human induced pluripotent stem cells (hiPSCs) were cultured in feeder-free monolayers on Geltrex (ThermoFisher Scientific, cat. no. A1413302) coated plates and fed daily with mTESR Plus (StemCell Technologies, cat. no. 85850). When 80-90% confluent, hiPSCs were dissociated using 0.5mM EDTA (Life Technologies, cat. no. 15575020) at 37°C for 5 minutes, EDTA aspirated, cells collected in mTSER Plus and plated onto Geltrex-coated plates at desired densities. All cells were maintained at 37°C and 5% CO_2_. A comprehensive list of all hiPSC lines used in this study can be found in supplementary key resources table.

Differentiation of *SNCA* A53T patient-derived hiPSCs into cortical neurons was performed using a modified, published protocol^82^. Briefly, hiPSC were cultured in Geltrex-coated 6-well plates until 100% confluent, media was then replaced with N2B27 with dual SMAD inhibitors: half DMEM/F12 (Gibco/ThermoFisher Scientific, cat. no. 10565018) and half Neurobasal-A (Gibco/ThermoFisher Scientific, cat. no. 12348017) as the base, 5µg/ml insulin solution human (Sigma, cat. no. I9278), 100µM 2-mercaptoethanol (Gibco/ThermoFisher Scientific, cat. no 21985-023), 1x MEM Non-essential amino acids (Gibco/ThermoFisher Scientific, at. no. 11140050), 0.5x N2 Supplement (Gibco/ThermoFisher Scientific, cat. no.

17502048), 1% GlutaMAX (Gibco/ThermoFisher Scientific, cat. no. 35050038), 1% penicillin-streptomycin (Gibco/ThermoFisher Scientific, cat. no. 15140122), 0.5x B27 Supplement (Gibco/ThermoFisher Scientific, cat. no. 17504044), plus Dual SMAD inhibitors 10µM SB431542 (Tocris, cat. no. 1614) and 1µM Dorsomorphin (Tocris, cat. no. 3093). The media was replaced every day for 10-12 days until a neuroepithelium monolayer appeared.

The neuroepithelium sheet was dissociated by adding 500µl of 10mg/ml Dispase directly to the well and incubating at 37°C for 10-15 minutes. The layer of cells was transferred to conical falcons containing DPBS and broken up into smaller clumps before replating in geltrex-coated plates in N2B27 at a ratio of 1:2. The media was then replaced every two days with fresh N2B27.

After 30 days of induction, cells were dissociated into single cells using Accutase (STEMCELL, cat. no. 7922) at 37°C for 5-10 minutes. Cells were collected in conical falcons, diluted 1:5 in DPBS and centrifuged at 300g for 5 minutes. Cells were resuspended in N2B27 media with ROCK inhibitor (Tocris, cat. no. 1254/50), counted and a cell suspension was prepared at a density of ∼4 x 10^5^ cells/ml before platting on Geltrex-coated plates. The media was replaced every 3-4 days and experiments performed at 60-100 days.

The CRISPRi-i3N iPSC line in the WTC11 background used in this study were engineered to express mNGN2 under a doxycycline-inducible system in the AAVS1 safe harbour locus and to constitutively express dead Cas9-KRAB transcriptional repressor fusion protein at the CLYBL promoter safe-harbour locus. This iPSC line was a kind gift from the laboratories of M.E. Ward and M. Kampmann, the generation of which has been previously described ^42,43^

For the neuronal differentiation of CRISPRi-i3N iPSCs, the protocol previously described by Tian et al. 2019 was followed with minor modifications^43^. In brief, at day 0, iPSCs were dissociated into single cells using Accutase as previously described. The supernatant was aspirated, and cells resuspended in Pre-Differentiation media containing the following: KnockOut DMEM/F12 (Gibco/ThermoFisher Scientific, cat. no. 12660-012) as the base, 1x MEM non-essential amino acids (Gibco/ThermoFisher Scientific, cat. no. 11140050), 1x N2 Supplement (Gibco/ThermoFisher Scientific, cat. no. 17502-048), 10ng/ml of NT-3 (PeproTech, cat. no. 450-03), 10ng/ml of BDNF (PeproTech, cat. no. 450-02), 1µg/ml of laminin (Merck, cat. no. L2020), 10nM ROCK inhibitor (Tocris, cat. no. 1254/50) and 2µg/ml of doxycycline (Sigma, D9891) to induce expression of mNGN2. iPSCs were counted and plated at 5 x 10^5^ cells per Geltrex-coated well of a 6-well plate in 2ml of Pre-Differentiation media, or at 4-5 x 10^6^ cells per Geltrex-coated T75 flask in 10ml of Pre-differentiation medium. The media was replaced every day for the following 2 days with Pre-differentiation media without ROCK inhibitor. On day 3, pre-differentiated cells were dissociated into single cells with Accutase, centrifuged as previously described, and pelleted cells were resuspended in Neuronal Media containing the following: modified N2B27 as the base using B27 Supplement without antioxidants (Gibco/thermoFisher Scientific, cat. no 10889038), 10ng/ml NT-3, 10ng/ml BDNF, 1µg/ml of laminin and 10nM ROCK inhibitor.

Pre-differentiated cells were counted, a cell suspension prepared at a density of 5 x 10^5^ cells/ml in Neuronal Media and cells plated by adding the following volumes to Poly-L-Ornithine (Sigma, P4957) and laminin coated plates: 140µl/well of a 96-well plate, 200µl/well of a 8-chamber ibidi, 500µl/well of a 24-well plate and 2ml/well of a 6-well plate. On Day 7, half of the media was removed, and an equal volume of fresh Neuronal Media without ROCK inhibitor was added. On day 14, half of the media was removed and twice that volume of fresh medium without ROCK inhibitor was added, on day 21, one-third of the media was removed and replaced with twice that volume of fresh media without ROCK inhibitor, and on day 28, one-third of the medium was removed and an equal volume of fresh medium was added. Experiments were performed on i3N neurons aged between 15-30 days.

Generation of the pLVX-EF1α-mitoSRAI-IRES-Puro plasmid and second-generation lentiviral particles have been previously described^83^. Stable expression of mitoSRAI was established through reverse transduction of 1×10^6^ i3N hiPSCs in mTeSER Plus medium supplemented with 5µg/ml polybrene (Sigma) and 10µM ROCK inhibitor. Stably expressing cells were selected with 1µg/ml puromycin (M P Biomedicals UK) for > 3 weeks prios to any i3N inductions.

### Microglia-like cell differentiation

iPSC microglia-like cells (iMGLs) were differentiated from *SNCA* A53T patient-derived and CRISPRi iPSCs using a modified version of published protocols^39,62^. In brief, to generate embryoid bodies at day 0, iPSCs cultured in mTSER Plus media until 60-80% confluency were dissociated to single cells with Accutase, collected and transferred into 5ml of DPBS in 15ml falcons. Cells were pelleted by centrifuging at 300g for 3 minutes and resuspended in Embryoid Body media containing: mTSER, 10µM ROCK inhibitor, 50ng/ml BMP-4 (Petrotech, cat. no. 120-05), 20ng/ml SCF (Peprotech, cat. no. 300-070), and 50ng/ml of VEGF (Peprotech, cat. no. 100-02). 10,000 cells were plated per well in 96-well ultra-low attachment plates in 100µl of embryoid body media, centrifuged at 800rpm for 3 minutes before transferring to a tissue culture incubator. Embryoid bodies were cultured for 4 days with a half media top-up at day 2. On day 4, embryoid bodies were carefully collected and transferred into a 15ml falcon and left to settle. The embryoid media was aspirated and 5ml of Myeloid Differentiation media added containing: X-VIVO 15 (Lonza, cat. no. BE02-060F) as the base, 1% GlutaMAX, 1% Pen/Strep, 2-mercaptoethanol, 100ng/ml of M-CSF (Peprotech, cat. no. 300-25) and 25ng/ml of IL-3 (Peprotech, cat. no. 200-03). Approximately 96 embryoid bodies were transferred into T75 flaks containing 10ml Myeloid differentiation media. Half media changes were performed every 3-4 days. From day 25 onwards, microglia progenitors were harvested weekly from the suspension and plated in microglia maturation containing: X-VIVO 15 as the base, 1% glutaMAX, 1% Pen/Strep and 100ng/ml of M-CSF. After platting, microglia were further matured for 7 days before using them for experiments. Phagocytosis ability was assessed by incubating iMGLs with fluorescently labelled beads (AF594 Beads) and imaged live after 24h using a Zeiss 880 confocal microscope with 40x, 1.4 N.A oil objective and a pinhole of one airy units (AU).

### CRISPR/Cas9 mediated *SNCA* A53T point mutation in CRISPRi-i3N iPSCs

The experimental procedure for generating *SNCA* A53T CRISPR clones in CRISPRi-i3Ns iPSCs was adapted from Skarnes et. al 2019 with minor alterations^84^. The synthetic sgRNA were purchased from Synthego (5’-GTGGTGCATGaTGTGGCAAC (AGG)) and were resuspended in TE buffer at a concentration of 4µg/ml. The Alt-R HDR modified ssODN purchased from IDT (5’-TACTTTAAATATCATCTTTGGATATAAGCACAATGGAGCTTACCTGTTG**t**CACACCAT GCACCACTCCCTCCTTGGTTTTGGAGCCTACAAAAACAAATT) were resuspended in DPBS at a concentration of 200pmol/ml. Prior to nucleofection, Cas9 RNPs were assembled by mixing 16µg of sgRNA and 20µg of Alt-R™ S.p. HiFi Cas9 Nuclease V3 (IDT, cat. no. 1081060) and incubated at RT for 30-45 minutes. Prior to nucleofection, 1µl of ssODN was added to the pre-assembled Cas9 RNP. To deliver the Cas9 RNP and oligo, 8 x 10^5^ iPSCs were resuspended in 100µl complete P3 solution and the assembled Cas9 RNP and cells were nucleofected using the Amaxa 4D Nucleofector (Lonza), the “Primary Call P3” program and the pulse code “CA-137”. Cells were then transferred to a 6-well plate and 30µM Alt-R HDR Enhancer V2 (IDT) was added to the cells prior to transferring to a 37°C/ 5% CO_2_ incubator for two days.

Once the iPSCs have reached 80% confluency, cells are dissociated into single cell with Accutase and 800 cells transferred to a geltrex-coated 10cm dish, and single cell colonies are allowed to grow for 8-10 days. Single cell-derived colonies were picked with the aid of a dissecting microscope and a p200 pipette and transferred into two 96-well plates in duplicate, one coated with geltrex and containing mTSER Plus to be kept in culture, and the other plate was used for DNA lysis, PCR amplification and Sanger sequencing using the following primers: SNCA_SF1: 5’-GTATTGAAAACTAGCTAATCAG, SNCA_SR1: 5’-ATGTTCTTAGAATGCTCAGTG. Successfully edited clones were amplified and banked following Karyostat verification.

### Immunocytochemistry

For immunocytochemistry (ICC), cells platted in 8-well chambers were washed with PBS and fixed with 4% paraformaldehyde for 15 minutes at room temperature. Following three PBS washes, cells were permeabilized and blocked with 0,1% Triton X-100 and 5% BSA in PBS for 1 hour at room temperature. Cells were then incubated with primary antibodies diluted in 5% BSA in PBS at 4°C overnight (see key resources table for details). The following day, primary antibodies were washed three times with PBS and incubated with secondary antibodies diluted 1:1000 in 5% BSA in PBS for 1 hour at room temperature. Cells were then washed twice with PBS, incubated with Hoechst 33342 (cat. no. 62249) for 10 minutes at room temperature and washed once more with PBS. Cells were then left in fluorescence mounting medium and stored at 4°C until imaged.

Cells were imaged using a Zeiss 880 confocal microscope with 40x, 1.4 N.A oil objective and a pinhole of one airy units (AU). Between 3 and 5 fields of view were acquired per condition, all with a Z stack consisting of 5-7 slices and displayed as maximum intensity projections.

For mitophagy measures, 5×10^4^ day 3 i3Ns were plated on Geltrex-coated 96-well Phenoplates (Revvity) and cultured until day 21. Cells were treated with 1µM O/A and then fixed with 4% formaldehyde (Rockland; 20 mins at room temperature). Cells were washed 3 times with PBS, before blocking and permeabilisation with 1x PBS supplemented with 10% foetal bovine serum (FBS; ThermoFisher) and 0.2% Triton-X-100 (BioWorld) for 1 hour at room temperature. Cells were the incubated with primary antibody in blocking buffer overnight at 4°C: rabbit anti-pUb(Ser65). Cells were then washed 3 times with PBS prior to incubation with fluorescently conjugated secondary antibody in blocking buffer for 1 hour at room temperature: goat anti-rabbit IgG Alexa Fluor 568 with DRAQ5 included to counterstain nuclei. Confocal images were captured using Opera Phenix High-Content Screening system and 63x water objective (Perkin Elmer, RRID: SCR_021100). Columbus image analysis software (Revvity) was used to analyse pUb integrated density and calculate the mitophagy index using TOLLES/YPet intensity ratio (T/Y > 1 classed as YPet^neg^ and

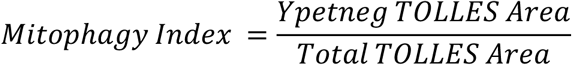

### Oxidative stress fluorescent live imaging (GSH, mROS, Het ROS)

To measure the antioxidant levels, reactive oxygen species (ROS) and mitochondrial ROS (mROS) generation, cells plated in 8-well chamber ibidis and were imaged live using either a confocal microscope (ZEISS LSM 880) with an illumination intensity limited to 0.1-0.2% laser output to prevent phototoxicity or a Nikon Ti2 inverted microscope. The levels of antioxidant were measured using a glutathione indicator, monochlorobimane (mCBl, ThermoFisherScientific, cat. no. M1381MP). Cells were incubated with 50µM mCBl in HBSS for 30 minutes at room temperature and cells imaged with a 405nm excitation laser and emission measured at 420-550nm.

mROS was assessed using MitoTracker Red CM-H2XRos dye (ThermoFisher Scientific, cat. no. M7513), a reduced non-fluorescent derivative of MitoTracker Red that fluoresces upon oxidation, accumulating in mitochondria. Cells were loaded with 2µM of MitoTracker Red CM-H2XRos for 5-7 minutes prior to imaging a time series with 5-10s intervals for 10 minutes with excitation at 560nm and emission above 600nm. The rate of mROS production was calculated as the slope of increase in fluorescence for each cell and normalised to WT or untreated cells.

To measure cytosolic ROS (mainly superoxide), cells were imaged on a Nikon Ti2 inverted microscope with Perfect Focus System, an ASI motorised XY stage with piezo Z and an Okolab environmental chamber with a CO_2_ mixer. Images were acquired using an Andor iXon Ultra897 EMCCD camera. Cells were excited with a Cairn FuraLED light engine optimised for 340 and 380 nm with a dichroic mirror T400lp (Chroma) and an emission filter ET510/80 m (Chroma), using a 40x 1.3 NA S Fluor objective. Cells were loaded with 4µM dihydroethidium (Het, ThermoFisher Scientific, cat. no. D11347) in HBSS for 2-3 minutes before acquiring a time series with 5-10s intervals for 7-10 minutes. The oxidised form (ethidium) was exited at 530nm and measured above 560nm and the reduced form was excited at 380nm and measured at 415-470nm. The ratio of the fluorescence intensity, resulting from its oxidised/reduced forms, was quantified and the rate of ROS production was determined by dividing the slope of the Het ratio against the basal gradient of WT cells.

### α-Synuclein intracellular Förster resonance energy transfer (FRET)

Fluorescently labelled monomeric A53T αsyn FRET pairs (500nM total: 250nM AF488 donor and 250nM AF594 acceptor) were added to cell media and incubated at 37°C for 48 hours. Prior to imaging, dyes and treatments were removed and cells washed in HBSS. In addition, a set of control wells were incubated for 48 hours with 500nM of AF488 donor only. Cells were imaged on a ZEISS LSM 880 confocal microscope, the donor αsyn was excited at 488nm and imaged at 493-566nm, the acceptor was excited at 594nm and imaged at 597-678nm, the FRET signal was recorded with excitation at 488nm and emission at 597-678nm.

### Lipid peroxidation detection

The levels of lipid peroxidation were assessed both by flow cytometry and high-content confocal live imaging with the C11-BODIPY 581/591 fluorescent probe (ThermoFisher Scientific, cat. no. D3861). For lipid peroxidation measurements by flow cytometry, cells were dissociated into single cells with Accutase, transferred to 1.5ml Eppendorf tubes, centrifuged at 300g for 5 minutes, resuspended in 100µl DPBS with LIVE/DEAD blue fluorescent probe (ThermoFisher Scientific, cat. no. L23105) to identify live and dead cells, and incubated for 15 minutes at 37°C. Then, an additional 100µl DPBS with double the concentration of C11-BODIPY 581/591 (4µM) was added to the cells and cells were incubated for are further 15 minutes at 37°C before analysing using the BioRad ZE5 cytometer. Data were collected from at least 10,000 live and single cells and C11-BODIPY mean intensity fluorescence (MIF) was measured in the living single cell population. Data was analysed using the FlowJo software, and lipid ROS levels were determined by the ratio of oxidised over reduced C11-BODIPY MIF and normalised to WT cells.

For high-throughput confocal imaging of lipid ROS, cells cultured in PLO and laminin coated 96-well plates were stained with 2µM C11-BODIPY 581/591 in HBSS for 15 minutes at 37°C. Then, the fluorescent probe was removed, and cells were treated with RSL3 +/-compounds in HBSS and cells were immediately imaged on the Operetta High content imaging system using the 488nm and 594nm channels, the confocal setting and 20x objective, every 30 minutes for 6 hours. A minimum of 9 fields of view and a Z-projection of 6 slices were acquired per well, and images displayed and analysed as a maximum projection.

The accompanying software, Harmony, was used to store and analyse acquired images with a pipeline design to detect all total cells (stained with the reduced C11-BODIPY dye) and the MIF of both the reduced and oxidised C11-BODIPY forms measured. Lipid ROS levels were determined by the ratio of oxidised over reduced C11-BODIPY MIF over time and normalised to t=0 hours.

### Image analysis

ICC images and live-cell imaging were analysed using Fiji ImageJ. For fluorescent intensity readouts, a threshold value was set using control/WT images. This was then used to measure fluorescent intensity, area and integrated density readouts for all images. To analyse cytosolic ROS and mROS production, individual cell bodies were selected as ROI and intensity measured using the Fiji Image J plugin “Time Series Analyzer V3” for every step of the time series. These values were used to generate traces and normalised to t=0 hours.

To measure FRET intensity the donor (donor_ex_ and donor_em_), acceptor (acceptor_ex_ and acceptor_em_) and FRET channels (donor_ex_ and acceptor_em_) were z-projected and background subtracted. A threshold was applied to the donor and acceptor channels and ROIs with signal above threshold in both channels were masked and used to measure intensity of the FRET channel and donor channel independently. FRET efficiency was calculated by the following equation: FRET intensity / (FRET intensity + donor intensity). As a control, efficiency in the absence of the acceptor dye was also calculated within donor positive ROIs. FRET intensity was averaged across FOV’s for statistical analysis. For each experiment, values were normalised to the average of all the WT/controls in the dataset. All graphs and traces were plotted in Prism v.10 (GraphPad).

### ELISA assay

To measure the concentration of oligomeric α-synuclein, cells were pre-treated with 50nM CU-12991 for 5 days, lysed mechanically in PBS and analysed using human α-synuclein oligomer (non A4 component of amyloid precursor) ELISA kit (Cusabio, cat. No. CSB-E18033h) according to the manufacturer’s instructions. The levels of intracellular αsyn was normalised to protein concentration measured by Pierce BCA assay (Thermo scientific, cat. No. 23228) following manufactures instructions.

### RNA extraction and qPCR

RNA was extracted from snap-frozen cell pellets using the Maxwell RSC simply RNA cells kit (Promega, cat. no. AS1390), and the accompanying Maxwell RSC 48 instrument, according to manufacturers instructions. Following extraction, 2µg of RNA was reversed transcribed into cDNA using the High Capacity cDNA Reverse Transcription Kit (ThermoFisher Scientific, cat. no. 4368814). The qPCR was performed using the Powerup SYBR Green Master mix for qPCR (ThermoFisher Scientific, cat. no. A25743) on the QuantStudio 7 Flex. The gene expression levels were normalised to the housekeeping gene *GAPDH* or *HNRN* following the delta-delta Ct method.

### Ferroptosis-related gene over-representation analysis

We utilised a list of ferroptosis-related genes (N=484) from FerrDb V.2^36^ using genes identified as either drivers, suppressors or markers of ferroptosis. Using this gene set we performed an ORA with differentially expressed genes from publicly available transcriptomic datasets. ORA was performed using enricher (clusterprofiler^85^) with our curated gene set as TERM2GENE and all detected genes from Evans 2025^39^ supplied as the enforced universe (N=19,708). To investigate if ferroptosis-related genes were enriched in a specific cell type we performed the same analysis with pseudo-bulked single cell transcriptomics from astrocytes, microglia and oligodendrocytes^60^.

### Bulk RNA-sequencing

Sequencing libraries were prepared using the Watchmaker rRNA and globin depletion kit, as per manufacturer’s instructions. RNA was sequenced on a NovaSeq X with an average depth of ∼27 million paired-end reads with a fragment length of 100 base pairs, per sample. Raw reads were processed using the NextFlow (v.4.04.2) nf-core pipeline (v.3.14.0)^86^ – briefly, samples underwent trimming of low-quality reads and adapters, prior to alignment against the human genome (v.GRCh38.95) using STAR and salmon. Differential gene expression analysis was performed using DESeq2^87^ (RStudio). Firstly, replicate samples were collapsed, and transcripts present in all samples (counts>1) subsetted for further analysis. Variance stabilisation was performed prior to principal component analysis (PCA, prcomp). DESeq was performed using the following design ∼ RIN_quality + genotype + treatment + genotype:treatment. RIN was included in the design as a categorical variable to account for varying RNA quality at input, as measured by TapeStation. Finally, log_2_ fold-changes were shrunk with the ashr algorithm^88^ to account for confounding effects of counts on fold-changes. To infer the biological significance of these changes, ORA was performed using up and downregulated genes independently via ClusterProfiler^85^ and ReactomePA^89^ with 5% FDR correction.

### Lipid extraction

Cell extracts were added to a 1.5 ml Sarstedt tube and 100 µl of extraction solution, consisting of 45% acetonitrile, 30% chloroform, 10% DMSO, 10% ethanol and 5% water including deuterium labelled internal standards, was added. The internal standards were Ceramide-d3 d18:1/18:0 (Matreya #2201), HexCer-d3 18:1/16:0 (Cayman #24621), HexSph-13C6 d18:1 (Cayman #23212), LPC-d7 18:1 (Avanti #791643C), LPA-d9 16:0 (Cayman #33479), PC-d7 15:0/18:1 (Avanti #791637) and PE-d7 15:0/18:1 (Avanti #791638). Cells and extraction solutions were sonicated for 10 minutes, followed by shaking for 10 minutes. Finally, the samples were centrifuged for 5 minutes in room temperature at 16,900 x g and 90µl of the supernatant was transferred to LC glass micro vials.

### Analysis of lipids by targeted LC-MS/MS and data integration

The samples were analysed using a triple quadrupole mass spectrometer (Waters TQ-S) equipped with an ESI source, coupled to a binary Waters Acquity liquid chromatographic separation system. Detection was performed in multiple reaction monitoring mode (MRM), where the transitions from precursor ions to class-specific fragment ions were monitored. To ensure that adequate analytical setups were employed for each of the compound classes, we utilised three separate LC methods with the column chemistries C8 or HSS T3, coupled with class-specific MRM methods. Data were acquired in MassLynx 4.1 and transformed into text files using the application MSConvert from the package ProteoWizard29. Peak picking and integrations were performed using an in-house application written in Python (available via the GitHub repository https://github.com/jchallqvist/mrmIntegrate) which rendered area under the curve by the trapezoidal integration method. Each analyte was thereafter normalised to an internal standard.

## Statistical analysis

Statistical tests were performed using unpaired t-tests, one-way ANOVA or two-way ANOVA corrected with Tukey’s/Dunnetts post hoc tests and linear and non-linear regression curve fittings were performed using Prism 10 GraphPad Software. Results are expressed as means ± standard error of the mean (SEM) and significant p values set at 0.05. Data points represent biological replicates either number of inductions or number of individual cells lines/clones, if not stated otherwise.

## Author contributions

Conceptualization: L.M.S., S.G. Methodology: L.M.S., H.L.C., J.E., A.G., K.M., J.H., K.C., D.S., B.O., S.P. Investigation: L.M.S., A.P., H.L.C., J.E. Analysis: L.M.S., S.B., H.L.C., J.E. Visualisation: L.M.S., H.L.C., J.E. Writing – original draft: L.M.S. Review & editing: H.L.C., S.G., J.E., S.P. Supervision: S.G. Funding: S.G., H.P-F.

## Conflict of interest

The authors declare no conflict of interest.

## Acknowledgements

The authors wish to thank the donors of fibroblasts, the Francis Crick Institute Flow cytometry, advanced light microscopy, High-throughput, genomics and bioinformatics STPs for their help and equipment in conducting and analysing the experiments. This research was funded in part by Aligning Science Across Parkinson’s [ASAP-000509]. K. Cosker and D. Soltic were supported by Eisai, working within the Eisai:UCL Therapeutic Innovation Group (TIG) with funding to H. Plun-Favreau. B. The authors wish to also thank Mr. Ben Powney, co-head of Biology at Eisai for his intellectual input. O’Callaghan was supported by “Guarantors of Brain Postdoctoral Non-Clinical Fellowship”.

---

**Supplementary figure 1 – Ai-ii.** Schematic of NGN2 and dCas9 insertions into AAVS1 and CLYBL safe harbour loci, respectively, present in the i3N model. **B.** Schematic of the differentiation protocol of CRISPRi-i3N iPSCs into neurons. **C.** Representative brightfield images of the differentiation process from iPSC (day 0), through NPCs (day 3), to neurons (day 8). **D.** Representative immunocytochemistry images stained for neuronal markers; Map2 (red), btubIII (purple), NeuN (green), in addition to Hoechst (blue), scale bar = 20mm. **Ei-ii.** qPCR revealing knockdown of *GCH1* and *PLA2G6* (via two independent sgRNAs) compared to non-targeting and untreated (iPSC: **p= 0.0037, ***p= 0.0007, d13 neurons: *p= 0.023, **p= 0.0024, Two-way ANOVA). **F.** Schematic of CRISPR/Cas9 editing pipeline to generate i3N lines harbouring an A53T mutation within *SNCA*. **G.** Sanger sequencing confirming successful edit. **H.** Representative ICC showing A53T lines differentiate into mature neurons (MAP2 red, a-synuclein green), scale bar = 20mm. **I.** qPCR reveals increase in *SNCA,* neuronal maturity markers (*MAP2*, *SYN1*, *NeuN)* and decrease in pluripotency markers (*SOX2*, *NANOG*) over time.

**Supplementary figure 2 –** Normal karyotypes were confirmed for the dCas9 parental line, two CRISPR WT clones, and four out of six heterozygous A53T CRISPR clones.

**Supplementary figure 3 – A.** Schematic experimental design for transcriptomic and lipidomic analyses on treated (+/-6 hour 300nM RSL3) i3Ns in the absence of antioxidants (-AO). **B.** Heat map of Log2 fold change of genes within enriched transcription factor signalling pathways after RSL3 treatment in A53T i3Ns. **C.** Representative ICC images of cortical neuronal markers - Ctip2 (red) and TBR1 (yellow) – and Hoechst (blue) from isogenic control and A53T patient iPSC-derived cortical neurons (day 71, scale bar = 50mm). **Di-ii.** C11 BODIPY581/591 time course (0-6 hours) of patient-derived iPSC cortical neurons after RSL3 treatment with varying doses (0.5mM, 1mM and 5mM). Representative images of WT and A53T treated with 0.5mM RSL3. (p<0.0001, Extra sum-of-squares F test). Non-oxidised BODIPY= red, oxidised= green.

**Supplementary figure 4 – Ai-ii.** Trace of average cytosolic ROS production in A53T +/-pre-treatment with 50nM CU-12991 for 5 days based on DHE fluorescence (non-significant, extra sum-of-squares F test) and quantification of cytosolic ROS production normalised to WT (dotted line) (non-significant p= 0.48, two-tailed unpaired Student’s t-test). **Bi-ii.** Quantification of mCB intensity normalised to WT (dotted line) in A53T +/-50nM CU-12991 (*p= 0.039, two-tailed Student’s t-test), and representative images of mCB staining via confocal microscopy (yellow). **C.** Histogram of FRET efficiency across all voxels in image for WT/A53T +/-50nM CU-12991 (p<0.0001, Extra sum-of-squares F test).

**Supplementary figure 5 – A.** Representative images of IBA1 (yellow) and CX3CR1 (magenta) expression in iMGLs derived from WT and A53T NGN2-iPSC cells (scale bar = 100mm). **B.** qPCR of microglial markers (*CX3CR1* and *P2YR12*) in iMGLs derived from WT and A53T NGN2-iPSC cells. **C.** Representative images showing phagocytosis of AF594 beads by A53T and WT iMGLs after 24h incubation (scale bars = 20mm). **D.** Intracellular iron levels in WT and A53T i3N-derived iMGLs as measured by ICP-MS. **E.** Lipid peroxidation measurements (oxidised/non-oxidised BODIPY) via high-throughput confocal live imaging over 6 hours in WT and A53T iPSC patient-derived astrocytes (non-significant, Extra sum-of-squares F test). **Fi-Fiii.** Heatmaps of log2FC significant genes leading to pathways enrichments: ferroptosis, ferrous iron binding and intracellular iron ion homeostasis from hiPSC patient-derived iMGLs Evans et al. (2025).

## Notes

### Competing Interest Statement

The authors have declared no competing interest.

