## Supplementary figures and images for "The *SNCA* A53T mutation sensitizes human neurons and microglia to ferroptosis"

### Supplementary Figure 1

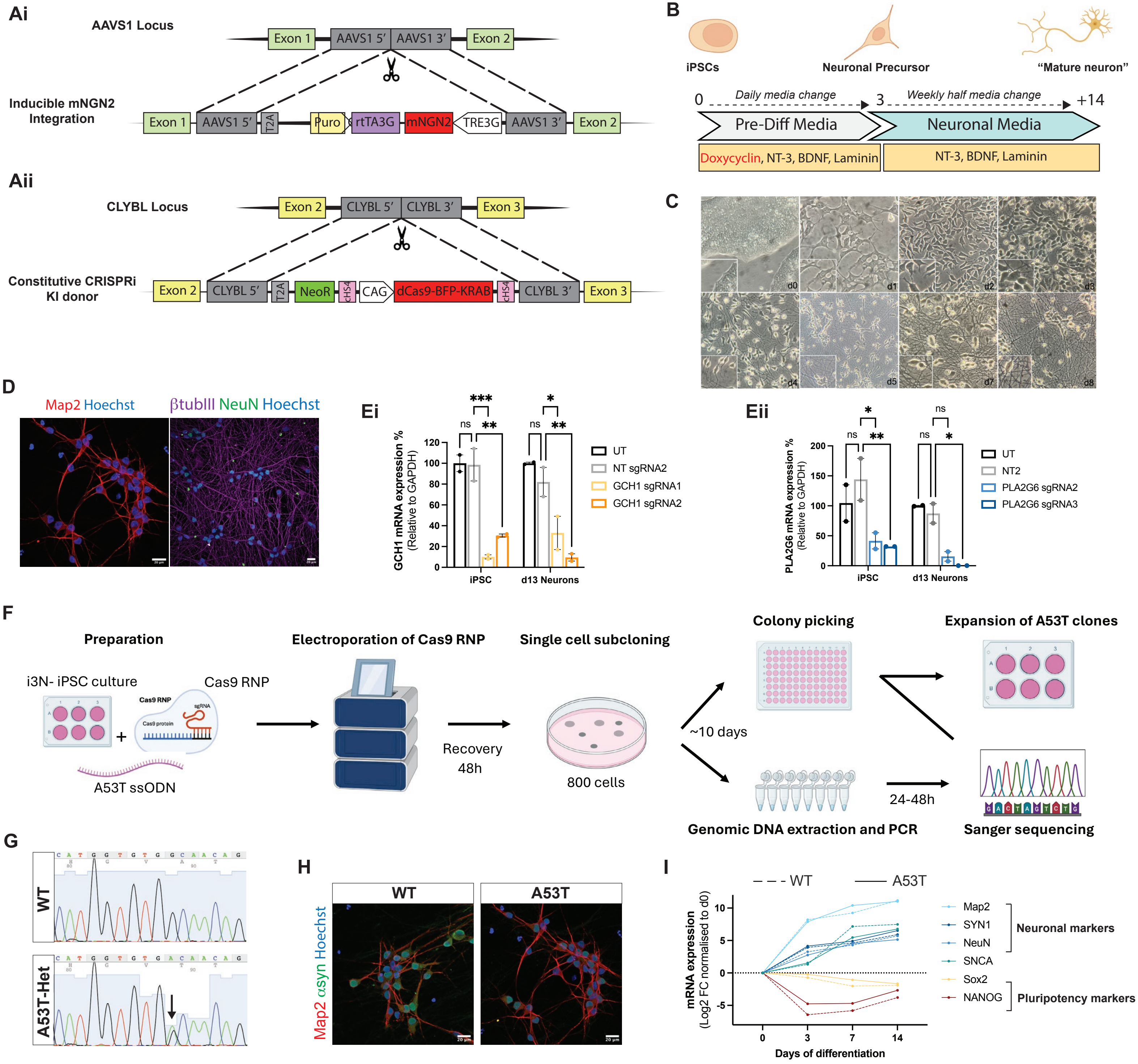

### Supplementary Figure 3

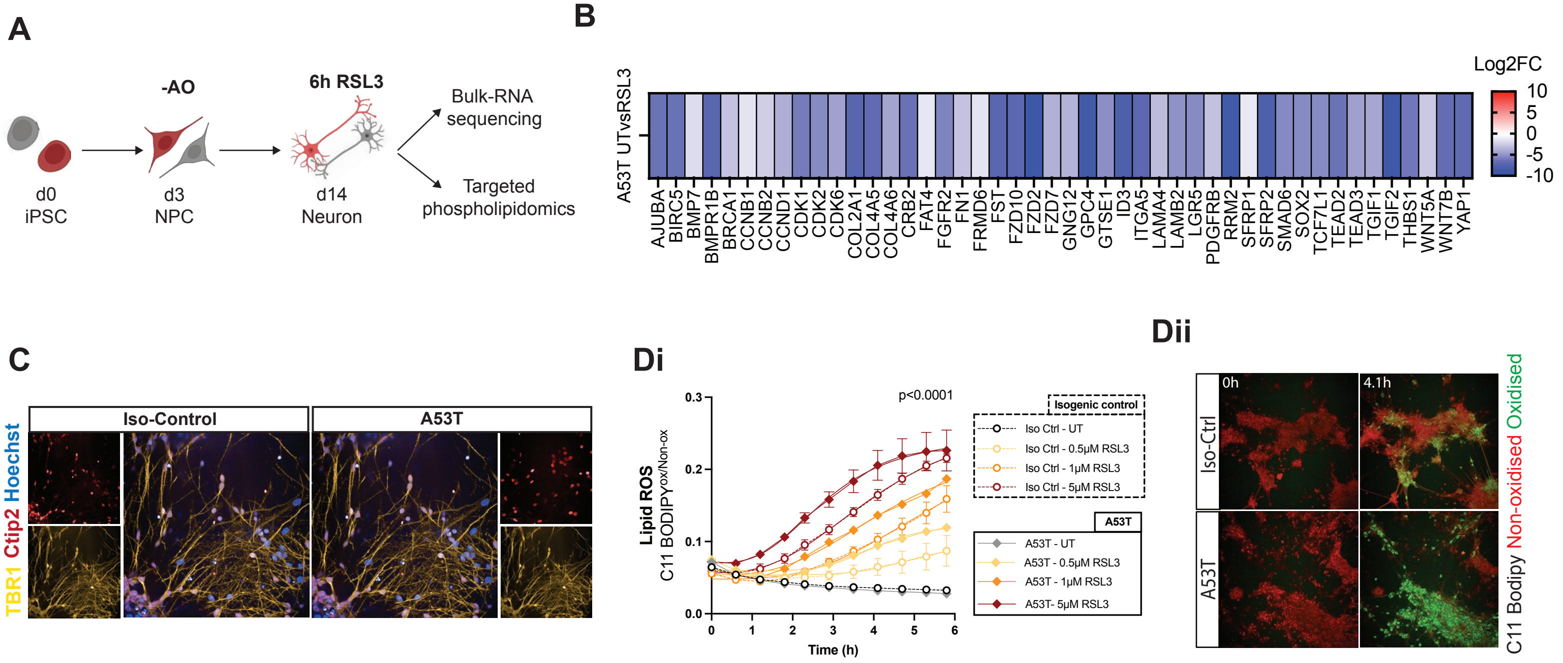

### Supplementary Figure 4

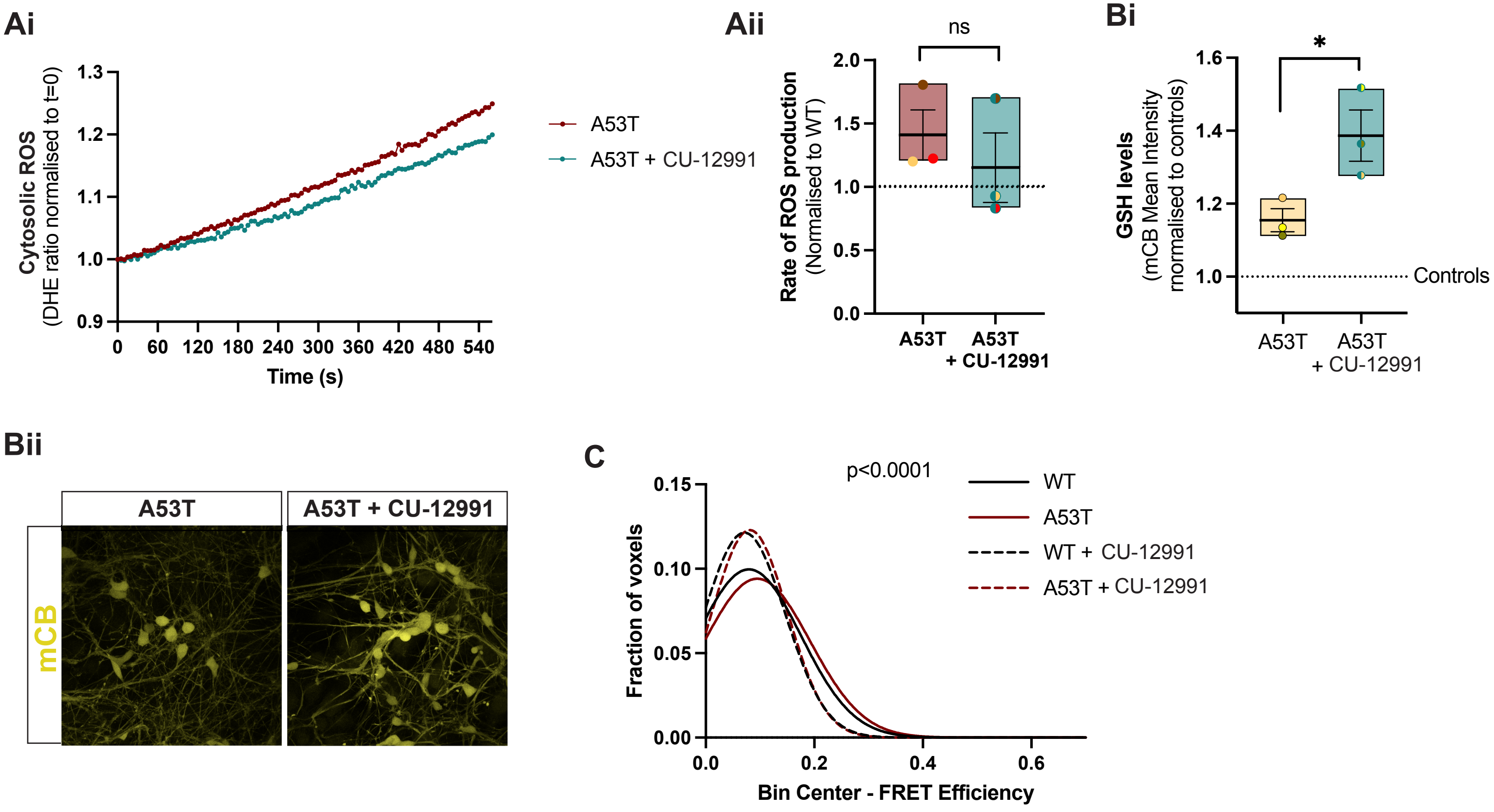

### Supplementary Figure 5

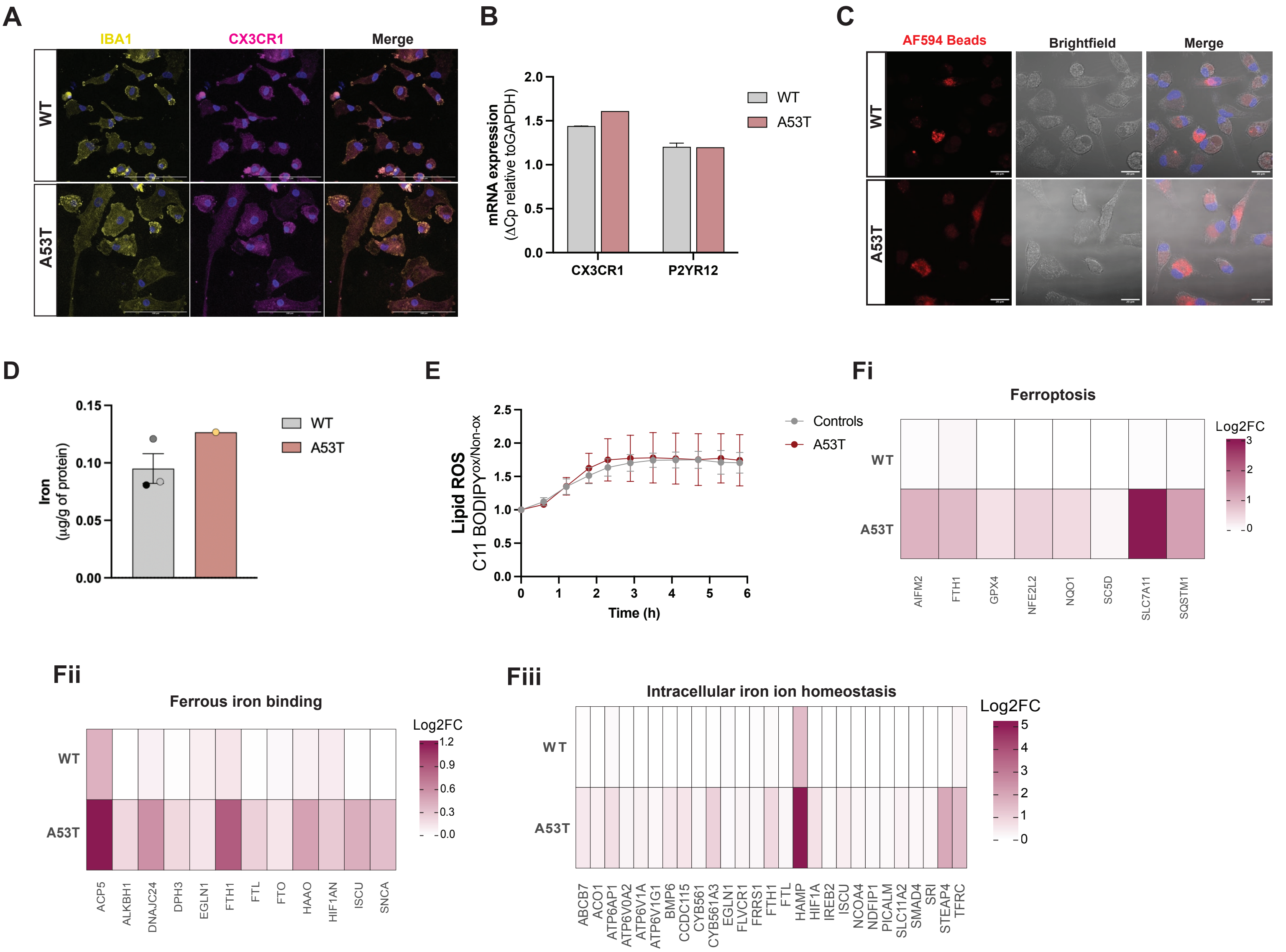
