## Supplementary Figure 2 for "The *SNCA* A53T mutation sensitizes human neurons and microglia to ferroptosis"

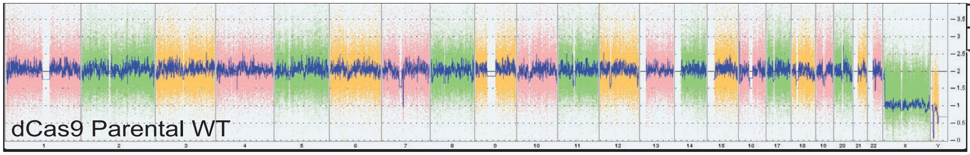


| Chromosome | Type | Cytoband start | Size (kbp) | CN |
|---|---|---|---|---|
| Y | Loss | p11.2 | 3,035 | 0 |

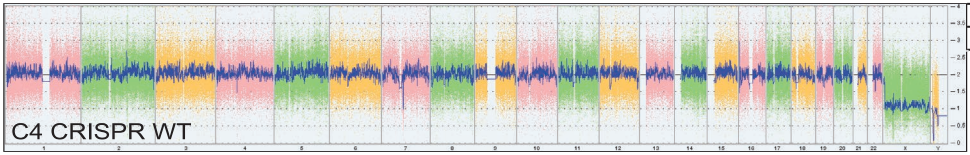


| Chromosome | Type | Cytoband start | Size (kbp) | CN |
|---|---|---|---|---|
| Y | Loss | p11.2 | 3,035 | 0 |

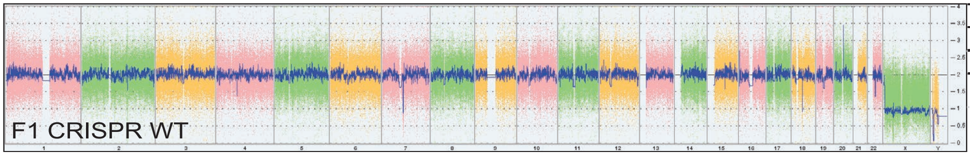


| Chromosome | Type | Cytoband start | Size (kbp) | CN |
|---|---|---|---|---|
| Y | Loss | p11.2 | 1,457 | 1 |
| Y | Loss | p11.2 | 3,044 | 0 |

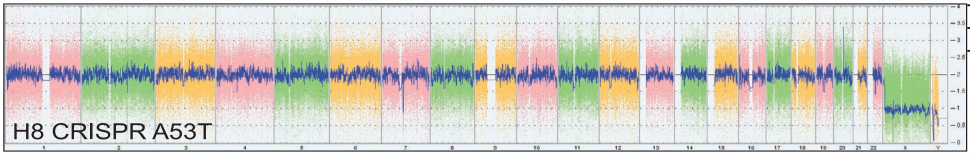


| Chromosome | Type | Cytoband start | Size (kbp) | CN |
|---|---|---|---|---|
| Y | Loss | p11.2 | 3,044 | 0 |

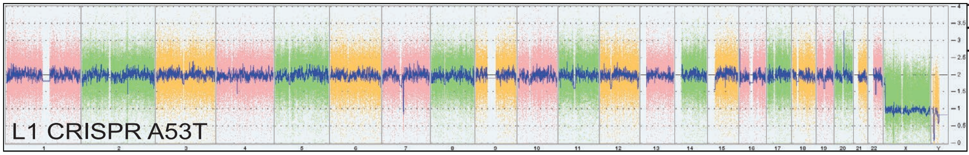


| Chromosome | Type | Cytoband start | Size (kbp) | CN |
|---|---|---|---|---|
| Y | Loss | p11.2 | 3,032 | 0 |

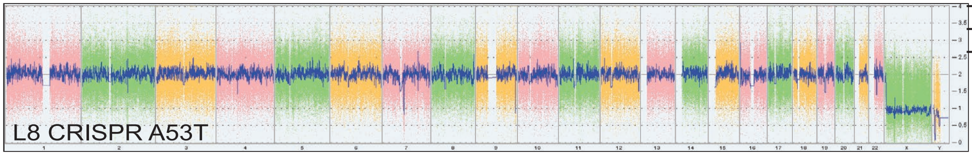


| Chromosome | Type | Cytoband start | Size (kbp) | CN |
|---|---|---|---|---|
| Y | Loss | p11.2 | 3,044 | 0 |

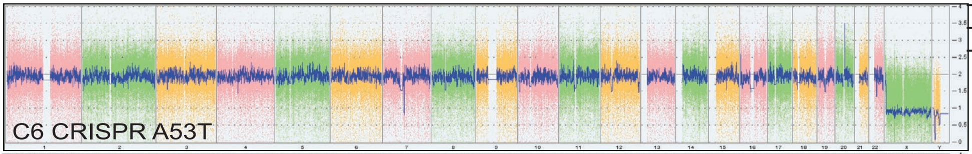


| Chromosome | Type | Cytoband start | Size (kbp) | CN |
|---|---|---|---|---|
| Y | Loss | p11.2 | 3,032 | 0 |

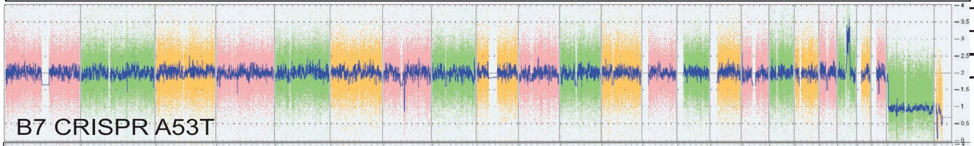


| Chromosome | Type | Cytoband start | Size (kbp) | CN |
|---|---|---|---|---|
| 20 | Gain | p11.2 | 13,096 | 3 |
| Y | Loss | p11.2 | 3,035 | 0 |

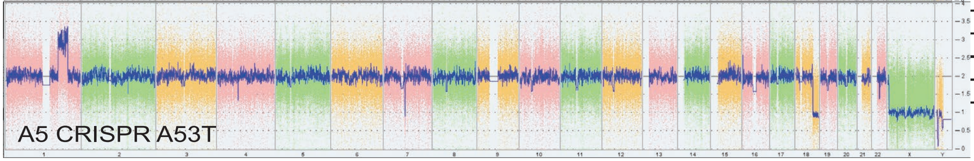


| Chromosome | Type | Cytoband start | Size (kbp) | CN |
|---|---|---|---|---|
| 1 | Gain | q35.1 | 32,794 | 3 |
| 18 | Loss | q21.31 | 21,864 | 1 |
| Y | Loss | p11.2 | 3,032 | 0 |
